# PRC2 disruption in cerebellar progenitors produces cerebellar hypoplasia and aberrant myoid differentiation without blocking medulloblastoma growth

**DOI:** 10.1101/2022.07.11.499567

**Authors:** Abigail H. Cleveland, Daniel Malawsky, Mehal Churiwal, Claudia Rodriguez, Frances Reed, Matthew Schniederjan, Jose E. Velazquez Vega, Ian Davis, Timothy R. Gershon

## Abstract

We show that the Polycomb Repressive Complex 2 (PRC2) maintains neural identity in Sonic Hedgehog (SHH)-driven cerebellar granule neuron progenitors (CGNPs) and SHH-driven medulloblastoma, a cancer of CGNPs. Proliferating CGNPs and medulloblastoma cells pass the neural fate commitment to their progeny through epigenetic mechanisms. The PRC2 mediates epigenetic regulation through histone methylation, and PRC2 inhibitors have been proposed for medulloblastoma therapy. We investigated PRC2 function in CGNPs and medulloblastoma by conditionally deleting PRC2 components *Eed* or *Ezh2* in CGNPs and analyzing cerebellar growth, cellular gene expression (scRNA-seq), and tumorigenesis in medulloblastoma-prone *Smo*-mutant mice. *Eed*-deleted CGNPs showed reduced growth, with decreased proliferation, increased apoptosis and inappropriate myoid differentiation. *Ezh2*-deleted CGNPs also showed myoid differentiation without reduced growth. *Eed*-deleted and *Ezh2*-deleted medulloblastomas similarly demonstrated myoid differentiation, but progressed more rapidly than PRC2-intact controls. The PRC2 thus maintained neural fate in CGNPs and medulloblastoma, but PRC2 disruption did not block SHH medulloblastoma progression.

## Introduction

During brain development, epigenetic inheritance specifies cell identities, directing progenitor cell differentiation along trajectories determined by their lineage. Thus, rhombic lip progenitors give rise to progeny with specific neural fates, including cerebellar granule neurons (CGNs) (1) and unipolar brush cell neurons (UBCs) (2). Lineage tracing by scRNA-seq shows that CGNPs differentiate into neurons and not into other types of cells (3). However, hyperactivation of SHH signaling can transform CGNPs, resulting in medulloblastoma (4–7), a primitive neuro-ectodermal tumor that is the most common malignant pediatric brain tumor. While most medulloblastoma cells adhere to a neural fate trajectory (8, 9), a small fraction of tumor cells differentiate as glia(3, 10). In giving rise to glia, SHH medulloblastoma cells demonstrates increased pluripotency beyond the neural fate trajectory of CGNPs while maintaining commitment to typically neuro-ectodermal fates. Epigenetic mechanisms thus maintain neuroectodermal lineage commitment through generations of proliferating progenitors during brain development and through generations of proliferating tumor cells in medulloblastoma.

PRC2 is a chromatin regulatory complex that has been shown to positively or negatively regulate neural progenitor proliferation in different contexts. The core subunits of the PRC2 include EZH1/2, EED, SUZ12, and RbAp46/48. By regulating trimethylation of H3K27 (H3K27me3), PRC2 suppresses CDKN2A, thus increasing neural progenitor proliferation in the hippocampus (11). However, PRC2 also inhibits SHH-induced transcriptional regulation by depositing H3K27me3 at bivalent chromatin sites in the promoter regions of SHH pathway target genes (12). As SHH signaling drives CGNP proliferation (13, 14) and medulloblastoma tumorigenesis (7, 15–18), PRC2-mediated repression of SHH target genes suggests the potential to inhibit proliferation in SHH-driven cells. In addition to affecting proliferation, PRC2 regulates hippocampal progenitor differentiation (11), consistent with of its role in fate commitment in diverse developmental contexts, from drosophila larva (19) to mammalian embryonic stem cells (20, 21). PRC2 may similarly contribute to cerebellar development by regulating CGNP proliferation and differentiation.

As in brain development, prior studies have shown divergent roles for the PRC2 in cancer, positively or negatively regulating tumor growth in different types of tumors (Deb et al., 2014; Scelfo et al., 2015). SHH medulloblastomas upregulate EZH2, the catalytic subunit of the PRC2, suggesting a growth-promoting function (12), and EZH2 inhibition has been proposed as a medulloblastoma therapy (22). A more complex relationship however, is suggested by the finding that pharmacologically increasing H3K27me3 levels in cultured medulloblastoma cells decreases tumor cell viability (12). The role of EZH2 in catalyzing inhibitory H3K27me3 marks and its non-canonical PRC2-independent functions has raised questions about the therapeutic potential of EZH2 inhibition (23–25).

To determine how PRC2 function affects cerebellar development and SHH medulloblastoma, we disrupted PRC2 activity in the CGNP-specific *Atoh1* lineage in genetically engineered mice through conditional deletion of *Eed* or *Ezh2*. We then bred the resulting PRC2-mutant mouse lines with mice genetically engineered to develop SHH medulloblastoma from CGNPs. Our data show that CGNPs and medulloblastoma cells require PRC2 to maintain neural fate commitment, and that EED is specifically required for cerebellar growth, but neither EED nor EZH2 are required for medulloblastoma progression.

## Results

### EED is required for proper cerebellar growth

CGNPs markedly increased H3K27 trimethylation as they differentiated in the early postnatal period. During the period of CGNP proliferation from postnatal day 1 (P1) through P15, cells from the CGNP lineage segregate spatially according to developmental state, with undifferentiated and early differentiating CGNPs in the external granule cell layer (EGL) and differentiated, post-mitotic CGNs in the internal granule cell layer (IGL). At P7, CGNs throughout the IGL expressed H3K27me3, while H3K27me3 was undetectable in the EGL (Fig. 1A). H3K27me3 increased in the CGN of the IGL by P21. H3K27me3 thus correlated with neuronal maturation, suggesting a role for the PRC2 in the differentiation process.

**Figure 1.**
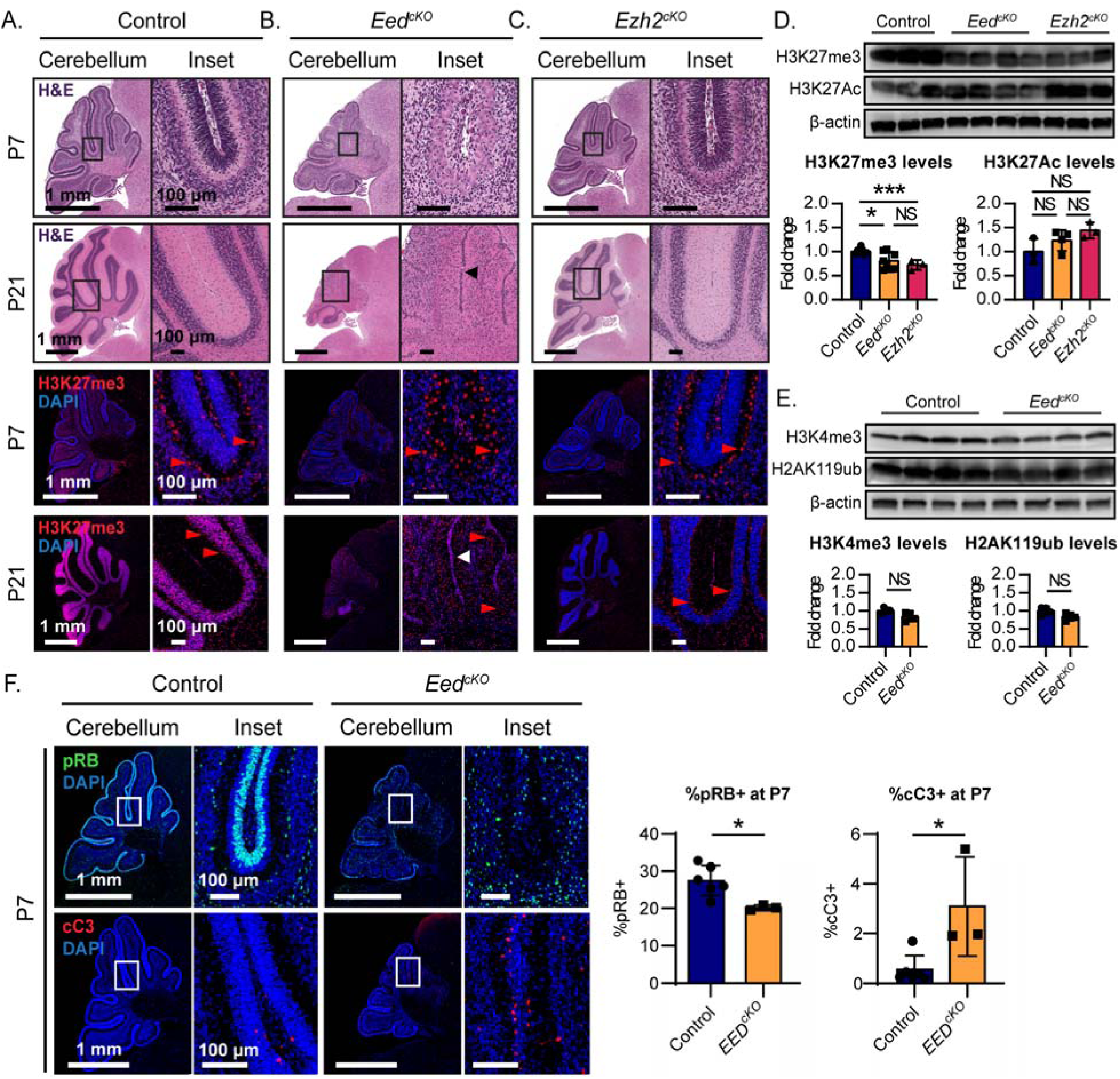
PRC2 function is developmentally regulated in the postnatal cerebellum and is required for normal development. **(A-C)** Representative sagittal sections of **(A)** control, (B) *Eed^cKO^*, and (C) *Ezh2^cKO^* cerebella at the indicated ages. H&E-stained sections show the distribution of cells in the EGL and IGL. IHC shows distribution of H3K27me3, with DAPI counterstain. Arrows indicate a residual EGL population in P21 *Eed* knockouts. **(D)** Western blot for H3K27me3 and H3K27Ac in replicate samples of indicated genotypes, quantified below. **(E)** Western blot for H3K4me3 and H2AK119ub in replicate samples of indicated genotypes, quantified below. **(F)** IHC for pRB and cC3 in representative sections, with quantitative analysis. *, **, and *** denote p<0.05, p<0.01 and p<0.001 respectively, relative to controls. Red arrowheads highlight Purkinje cells, identified by their typical large nuclei.

In light these data, we analyzed PRC2 function in CGNP using conditional genetic deletion. Prior studies showed that conditional *Ezh2* deletion in the dorsal neural tube at embryonic day E12.5 results in cerebellar hypoplasia, with decreased numbers of both Purkinje cells and CGNPs (26). However, the loss of Purkinje neurons may indirectly alter CGNPs, which depend on SHH released by Purkinje cells in order to proliferate (27). To disrupt PRC2 specifically in CGNPs, we deleted PRC2 genes conditionally by expressing *Cre* from the *Atoh1* (aka *Math1*) promoter which in the cerebellum is CGNP specific (28). We interbred *Math1-Cre* transgenic mice that express Cre recombinase in the CGNP population with mice harboring conditional alleles of either *Eed* (*Eed^fl/fl^*) or *Ezh2* (*Ezh2^fl/fl^*), to generate *Math1-Cre/Eed^f/f^* (*Eed^cKO^*) and *Math1-Cre/Ezh2^f/f^*(*Ezh2^cKO^*) mice.

*Eed* deletion resulted in marked depletion of the CGNP lineage. In P7 *Eed^cKO^* cerebella, the IGL of was sparsely populated with CGNs that were H3K27me3-, consistent with PRC2 disruption (Fig. 1B). In contrast, Purkinje cells remained H3K27me3+; these cells, which typically localize at the outer margin of the CGNs, were scattered throughout the depopulated IGL and were identified by their typical morphology with large nuclei and cell bodies (Fig.1B, red arrowheads). By P21, *Eed^cKO^* cerebella were clearly smaller than controls, with no clear IGL, and with an inappropriately persistent, small population of H3K27me3+ cells in the EGL (Fig.1B; arrow). Consistent with cerebellar impairment, *Eed^cKO^* showed tremor and ataxia and frequently fell while walking. EED was thus required for H3K27 tri-methylation in CGNPs and for cerebellar growth and function.

*Ezh2* deletion similarly reduced H3K27me3, but did not cause neurologic abnormalities or cerebellar hypoplasia (Fig. 1C). Thus while H3K27me3 was altered in both *Eed^cKO^* and *Ezh2^cKO^* mice, only *Eed* deletion resulted in cerebellar growth failure.

To determine if deletion of *Eed* or *Ezh2* produced different changes in chromatin regulation, we analyzed chromatin marks in *Eed^cKO^*and *Ezh2^cKO^* cerebellar lysates. Western blot analysis showed that *Eed* and *Ezh2* deletions similarly disrupted PRC2 function, and did not identify differences in the impacts on chromatin marks. Both deletions reduced H3K27me3 as expected (Fig. 1D). However, it was not possible to compare the extent of H3K27me3 suppression in *Eed*-deleted CGNPs vs *Ezh2*-deleted CGNPs, due to residual H3K27me3 from cerebellar cells outside the *Atoh1* lineage that were not subject to conditional deletion. We noted a trend toward increased H3K27 acetylation in both *Eed^cKO^*and *Ezh2^cKO^* (Fig. 1D) that was consistent with the previously observed increased H3K27Ac in H3K27me3-depleted embryonic stem cells (29, 30). *Eed^cKO^* CGNPS did not show differences in H2AK119 monoubiquitination, indicating that PRC1 function was not altered, and did not show differences in H3K4me3 (Fig. 1E). Within the resolution of our western blot assays, therefore, deletion of *Eed* or *Ezh2* resulted in similar chromatin changes, limited to H3K27 modification, suggesting that both deletions primarily altered PRC2 function, rather than other aspects of CGNP chromatin regulation.

### *Eed* deletion decreases progenitor proliferation and increases apoptosis

We investigated developmental changes in *Eed^cKO^* CGNPs, which showed the more abnormal phenotype. We quantified expression of the proliferation marker phosphorylated-RB (pRB) and the apoptotic marker cleaved Caspase-3 (cC3), as both decreased CGNP proliferation and increased CGNP apoptosis can cause cerebellar hypoplasia (13, 31–33). *Eed* deletion was previously shown to decrease proliferation and increased apoptosis in hippocampal progenitors (11). Similarly, *Eed^cKO^*CGNPs showed decreased proliferation and increased apoptosis (Fig. 1F), implicating both processes in the cerebellar hypoplasia of *Eed^cKO^*mice.

### *Eed* deletion produces discrete patterns of transcriptomic change

To identify transcriptional patterns specific to *Eed-*deleted cells, we subjected *Eed*-deleted and control CGNPs to scRNA-seq analysis. We harvested cerebella from 3 replicate P7 *Eed^cKO^*mice and subjected them to Drop-seq bead-based scRNA-seq preparation (Macosko et al., 2015). We then sequenced the resulting bar-coded libraries and identified cells by bead-specific bar codes. After QC and filtering, we included 2377 cells for analysis. We compared these cells to previously sequenced data from 6631 cells from 5 replicate P7 WT cerebella. To adjust for differences in sequencing depth in *Eed^cKO^*and WT cells, we down-sampled the WT cells to 46.5% of their original depth to achieve similar sequencing depth between conditions, consistent with best practices (34).

We subjected scRNA-seq data from *Eed^cKO^* and WT cells to principal component analysis (PCA) and Louvain clustering, as in our prior studies (3, 35, 36), to identify 20 clusters, numbered from 0, most populous, to 19, least populous (Supplementary Fig.1). We then determined cluster-specific gene expression profiles by comparing the expression of each detected gene in cells within the cluster versus all cells outside the cluster (Supplementary Data 1). In Clusters 4 and 6, we noted that cluster-specific genes were expressed by discrete subpopulations, suggesting that further sub-clustering of these clusters would be informative. Re-iterative clustering split Cluster 4 into 4_0 and 4_1 and Cluster 6 into 6_0 and 6_1, which showed discrete, cluster-specific patterns of gene expression (Supplementary Data 1). We mapped these 22 clusters by color code on the UMAP projection to visualize the different populations (Fig. 2A).

**Figure 2.**
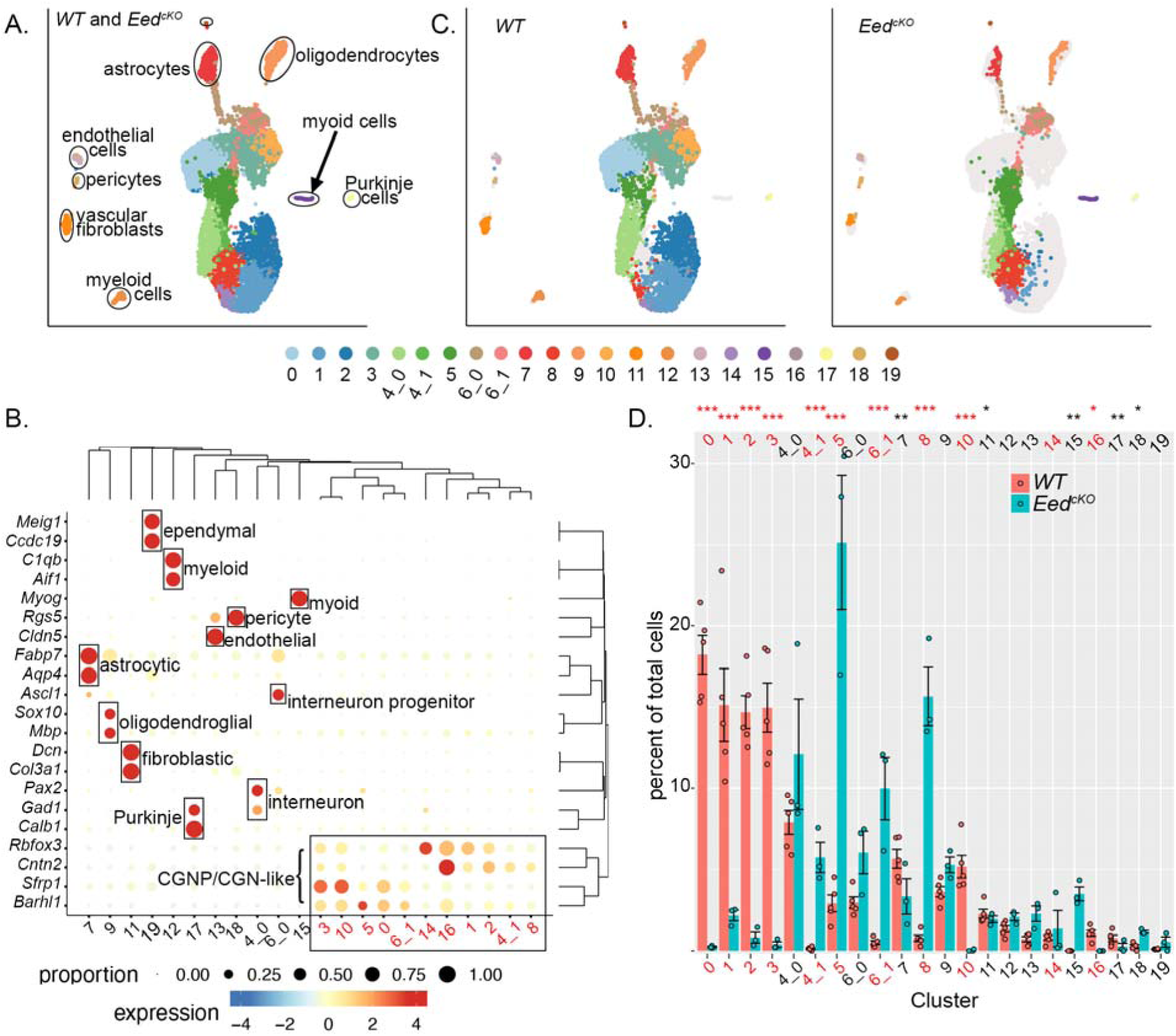
scRNA-seq shows differences in the composition of *Eed^cKO^* cerebella, including a population of myoid cells not found in normal brain. (**A**) UMAP projection of cells from *Eed^cKO^* and control cerebella, grouped by transcriptomic similarities into color-coded clusters. **(B)** Bubble Plot shows the magnitude and frequency of the expression of indicated cell type markers in each cluster. The genes and clusters are ordered according to the hierarchical clustering analysis, with groupings indicated along the top and right margins of the Bubble Plot. **(C)** UMAP projection of cells from *Eed^cKO^* and control cerebella, disaggregated by genotype, with clusters indicated by the same color code as in (A). **(D)** Comparison of each normalized cluster population in *Eed^cKO^*and control cerebella. Dots represent values for individual replicates, bars indicate the means and whiskers indicate the SEM. *** indicates p<0.0001,** indicates p<0.001, * indicates p<0.05; Dirichlet regression was used to compare normalized cluster populations. In **(B)** and **(D)**, CGNP and CGN cluster numbers are presented in red for clarity.

The differential gene expression patterns identified the cell-type of each cluster and demonstrated diverse cells typical of cerebellar tissue (Fig. 2A,B; Table 1). Markers *Sfrp1*, *Barhl1*, *Cntn2* and *Rbfox3* identified CGNP lineage cells in a spectrum of differentiation states, from cycling CGNPs to differentiated CGNs (Clusters 0, 1, 2, 3, 4_1, 5, 6_1, 10, 14, 16, 17). Different markers identified diverse types of non-neural cells, including astrocytes (Cluster 7), oligodendrocytes (Cluster 9), vascular fibroblasts (Cluster 11), myeloid cells (Cluster 12), endothelial cells (Cluster 13), pericytes (Cluster 18), and ependymal cells (Cluster 19). Markers also identified neural populations that were outside the *Atoh* lineage but expected in the cerebellum, including gabaergic neural progenitors (Cluster 6_0), identified by expression of *Ascl1* and *Pax3*, gabaergic interneurons (Cluster 4_0) identified by *Pax2* and *Gad1*, and Purkinje cells identified by *Gad1* and *Calb1* (3, 37). In contrast to all other clusters, which demonstrated markers expected in the brain, Cluster 15 expressed the muscle cell marker *Myog*, suggesting myoid differentiation.

**Table 1.** List of cell types identified via scRNA-seq. Cluster numbers and corresponding cell types, with *Atoh1*-lineage cells identified, CGNP and CGN clusters are in red, and population differences noted between *Eed^cKO^* and WT, with p values were determined by Dirichlet regression analysis.

| Cluster # | Designation | *Atoh1*-lineage? | Population difference | p value KO vs Control |
| --- | --- | --- | --- | --- |
| 0 | proliferating CGNPs | Yes | up in control | 1.00E-14 |
| 1 | differentiating CGNs | Yes | up in control | 8.40E-14 |
| 2 | early differentiating CGNPs | Yes | up in control | 1.00E-14 |
| 3 | late mitotic CGNPs | Yes | up in control | 1.00E-14 |
| 4_0 | interneurons | No |  |  |
| 4_1 | *Hox+* CGNPs | Yes | up in *Eed*<sup>CKO</sup> | 1.84E-05 |
| 5 | Proliferating CGNPs | Yes | up in *Eed*<sup>CKO</sup> | 1.03E-07 |
| 6_0 | early interneuron progenitors | No |  |  |
| 6_1 | *Cdkn2a+* proliferating CGNPs | Yes | up in *Eed*<sup>CKO</sup> | 3.34E-07 |
| 7 | astrocytes | No | up in control | 0.000213 |
| 8 | *Hox+* differentiating CGNs | Yes | up in *Eed*<sup>CKO</sup> | 2.91E-10 |
| 9 | oligodendrocytes | No |  |  |
| 10 | mitotic CGNPs | Yes | up in control | 5.67E-10 |
| 11 | vascular fibroblasts | No | up in control | 0.0351 |
| 12 | myeloid | No |  |  |
| 13 | endothelial cells | No |  |  |
| 14 | CGNs | Yes |  |  |
| 15 | myocytic cells | Yes | up in *Eed*<sup>CKO</sup> | 0.000945 |
| 16 | early differentiating CGNPs | Yes | up in control | 0.00127 |
| 17 | Purkinje cells | No | up in control | 0.0214 |
| 18 | pericytes | No | up in *Eed*<sup>CKO</sup> | 0.00831 |
| 19 | ependymal cells | No |  |  |

We dissaggregated the UMAP into separate projections from *Eed^cKO^*and WT mice, which demonstrated that each genotype distributed differently across the clusters (Fig. 2C). To compare the cluster populations in *Eed^cKO^* and WT samples statistically, since each replicate contributed different numbers of cells to the analysis, we normalized the population of cells in each cluster from each replicate to the total number of cells from that replicate. These proportional populations were inter-related by the normalization to the whole, and therefore could not be analyzed by individual t-tests, which assume independence. We therefore used Dirichelet regression analysis to compare the proportional cluster populations (38).

As suggested by the disaggregated UMAP (Fig. 2C), Dirichelet regression analysis showed that the populations of specific clusters were significantly different in *Eed^cKO^* cerebella compared to controls (Fig. 2D). We found significant differences in the populations of *Atoh1*-lineage cell types that were subject to conditional deletion, and also of clusters outside the *Atoh1* lineage. Within the *Atoh1* lineage, CGNP/CGN Clusters 0, 1, 2, 3, 10 and 16 were more populous in controls, while CGNP/CGN Clusters 4_1, 5, 6_1 and 8 were more populous in the *Eed^cKO^* (Fig. 2D). Outside the *Atoh1* lineage, Purkinje cells, astrocytes and vascular fibroblasts were increased in controls and pericytes were increased in the *Eed^cKO^*. These cell types were not subject to *Eed* deletion, and differences in their popluations reflect non-cell autonomous effects. Additionally, the myoid Cluster 15 was present only in the *Eed^cKO^* cerebella (Fig. 2C,D).

### *Eed* deletion permits divergence from the neural fate of CGNPs

In addition to *Myog*, Cluster 15 cells expressed multiple genes typical of muscle cells, including troponins, *Myl1*, *Cav3*, and *Smyd1* (Fig. 3A). This myoid differentiation was a specific effect of *Eed* deletion as we did not observe this myoid transcriptomic pattern in any cells from control mice. Cluster 15 cells also up-regulated *Cdkn1a* and *Cdkn1c* (Fig. 3A,B), which are known to be suppressed by the PRC2 in other types of cells (39–42). Based on the up-regulation of PRC2-suppressed genes, we infer that Cluster 15 cells were *Atoh1*-lineage cells with conditional *Eed* deletion, which resulted in a differentiation pattern that diverged from the expected CGNP trajectory.

**Figure 3.**
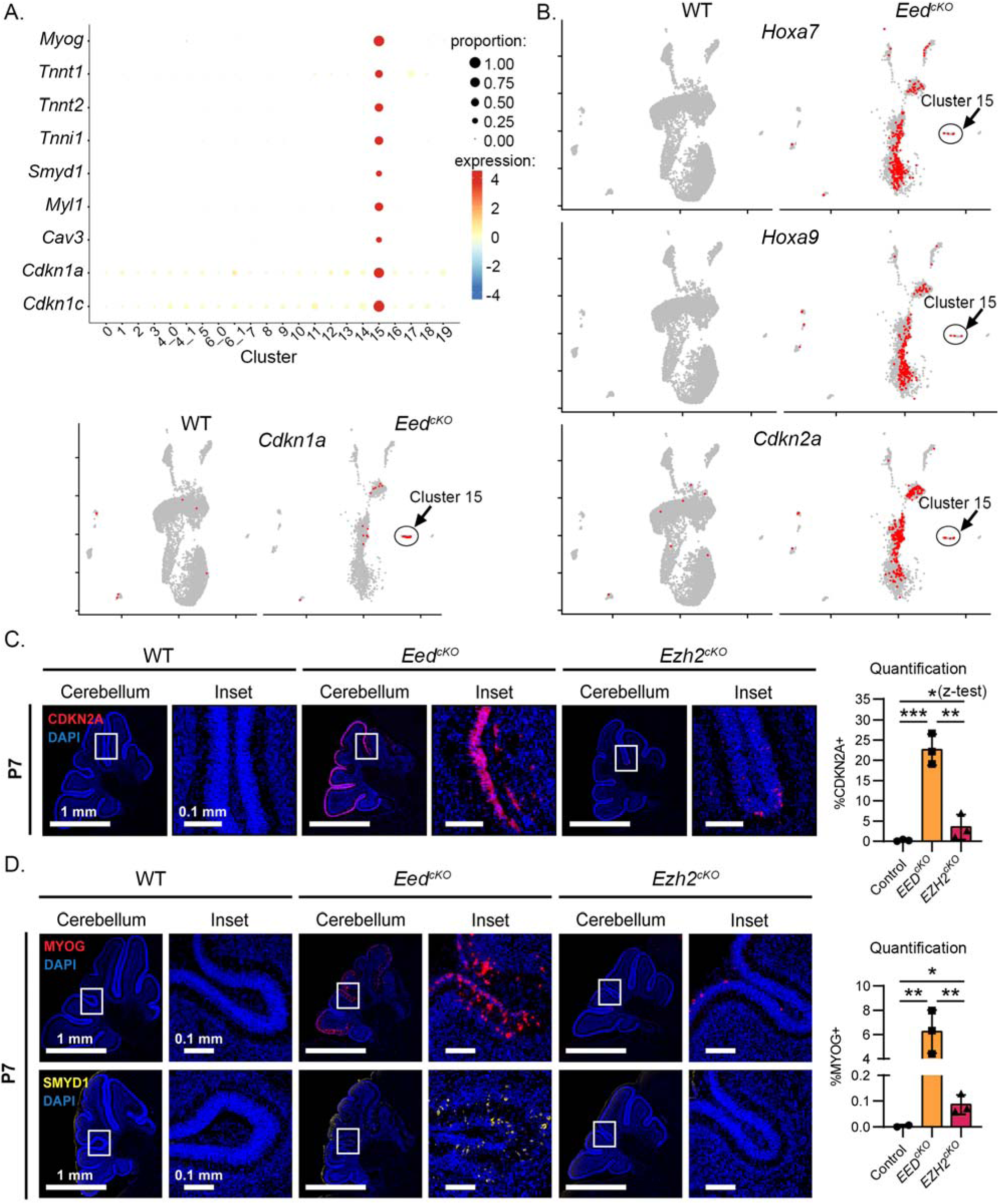
Expression of PRC2 targets in *Eed^cKO^* and *Ezh2^cKO^* cerebella. **(A)** Bubble Plot shows the magnitude and frequency of the expression of indicated genes in each cluster, and feature plot of *Cdkn1a*+ cells (red), color-coded over the UMAP from (2C). **(B)** Feature plots of *Hoxa9^+^*, *Hoxa7^+^*, and *Cdkn2a^+^*cells (red), color-coded over the UMAP from (2C). **(C,D)** Representative immunofluorescence in sagittal sections of WT, *Eed^cKO^*, and *Ezh2^cKO^*cerebella, showing expression of **(C)** CDKN2A or **(D)** MYOG and SMYD1, with DAPI counterstain. The fractions of cells in the cerebellum expressing each marker in 3 replicate stained sections are graphed on the right. Replicates of each genotype are compared by one-tailed Student’s t-test, except as noted. *, **, and *** denote p<0.05, p<0.01 and p<0.001 respectively.

### *Eed* deletion permits expression of PRC2-inhibited CDK inhibitors and *Hox* genes

While only Cluster 15 cells showed *Cdkn1a* and *Cdkn1c*, the broad set of *Atoh1*-lineage cells, including CGNPs, CGNs and Cluster 15 cells expressed other genes typically suppressed by the PRC2, including *Cdkn2a, Hoxa9*, and *Hoxa7* (42–45) (Fig. 3B). indeed, these markers differentiated the CGNP/CGN clusters enriched in *Eed^cko^* cerebella (Clusters 4_1, 5, 6_1 and 8) from the CGNP/CGN clusters enriched in WT samples (Clusters 0, 1, 2, 3, 10 and 16). The absence of *Cdkn2a, Hoxa9*, and *Hoxa7* in WT cells (Fig. 3B) indicates that each of these genes are normally suppressed by the PRC2 in CGNP-lineage cells.

### *Ezh2* deletion, like *Eed* deletion, permits myoid differentiation

We used immunohistochemistry (IHC) to determine if specific transcripts from PRC2-target genes were translated in *Eed^cKO^* cerebella, and to probe *Ezh2^cKO^* cerebella for similar patterns of expression. *Eed^cKO^* cerebella showed CDKN2A+ CGNPs throughout the EGL expressed (Fig. 3C). *Ezh2^cKO^* cerebella showed significantly smaller fractions of CDKN2A+ CGNPs (Fig. 3C). While the fractions of CDKN2A+ cells in *Ezh2^cKO^* cerebella were highly variable and not statistically significant when compared to WT by parametric testing, comparing the number of replicates with CDKN2A+ CGNPs in *Ezh^2cko^* versus WT showed p=0.05 (one-tailed 2 proportion z-test). Both PRC2 component mutations thus resulted in up-regulation of CDKN2A, with *Eed^cKO^* showing a higher fraction of affected cells.

Similarly, both *Eed^ckO^* and *Ezh2^ckO^*cerebella showed cells expressing myoid genes, with *Eed* deletion affecting more cells. While few cells expressed MYOG or SMYD1 in WT cerebella, *Eed^cKO^*cerebella showed numerous CGNPs expressing MYOG and SMYD1 (Fig. 3D). *Ezh2^cKO^*cerebella showed a smaller increase MYOG+ CGNPs, and no SMYD1+ CGNPs (Fig. 3D). Transcripts typically suppressed by the PRC2 in WT CGNPs, including *Cdkn2a* and myoid genes, were thus translated into proteins in both *Eed^ckO^* and *Ezh2^ckO^* cerebella. *Ezh2* deletion affected altered gene expression in fewer cells, which may explain the absence of a robust developmental phenotype.

### Increased activation of the intrinsic apoptotic pathway in *Eed^cKO^* CGNPs

To probe the mechanisms of increased cell death in *Eed^cKO^*CGNPs, we analyzed p53 function and the expression of genes regulating apoptosis. CGNPs are highly sensitive to p53-mediated activation of the intrinsic apoptotic pathway (46, 47) and also to direct activation the intrinsic apoptotic pathway triggered by changes in apoptotic regulators (33, 48). *Cdkn1a* expression in Cluster 15 suggested that *Eed* deletion might activate p53-dependent transcription, potentially mediating the observed increased apoptosis. To determine if other apoptotic regulators were altered in *Eed^cKO^* CGNPs, we compared the expression of pro-apoptotic BH3-only genes in the CGNP and CGN clusters of *Eed^cKO^* and control mice (Fig. 4A).

**Figure 4.**
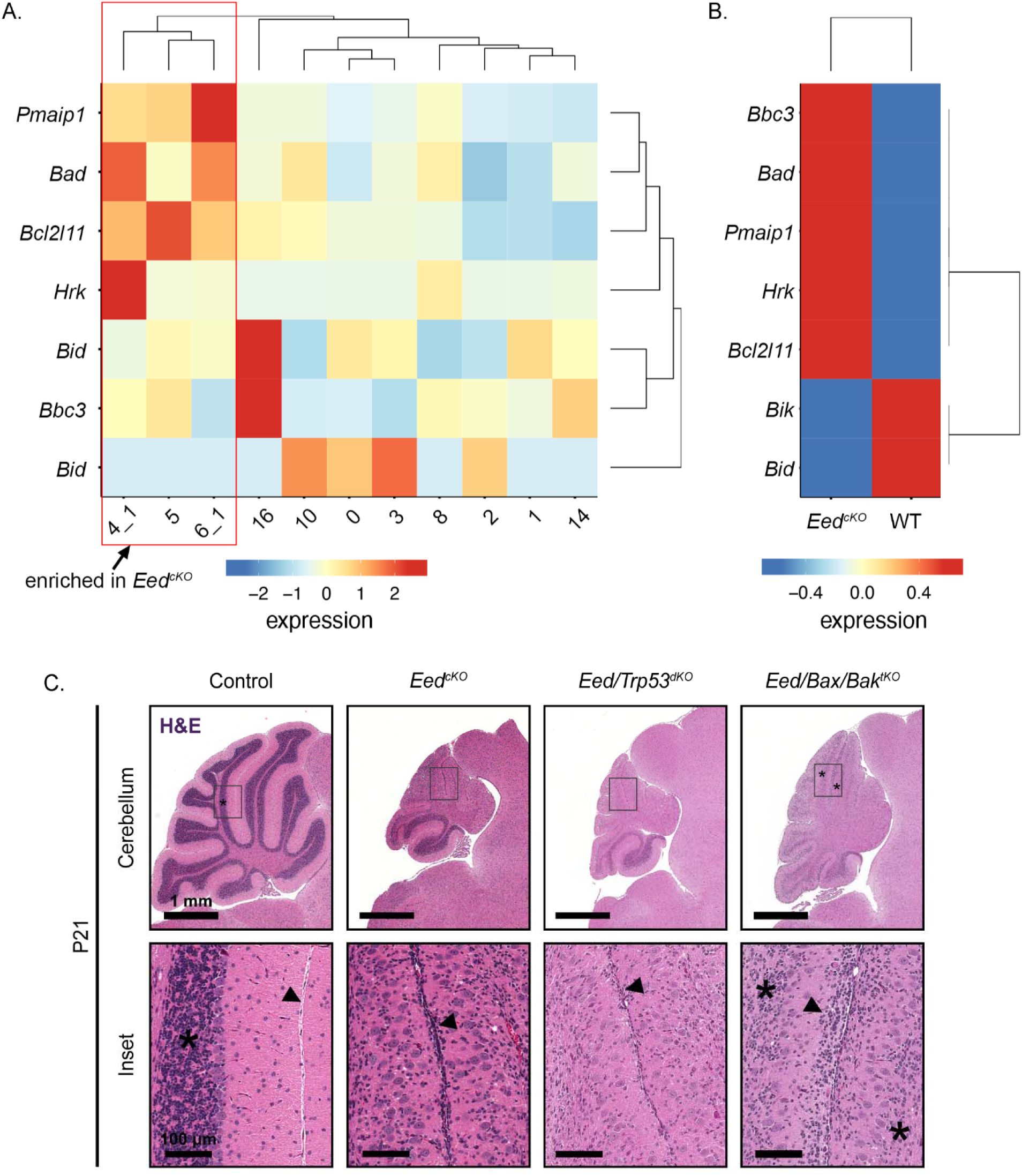
Increased spontaneous, p53-indepenedent CGNP apoptosis contributes to growth failure in *Eed*-deleted cerebella. (**A**) Heat map of BH3-only gene expression in each CGNP/CGN cluster. Hierarchical clustering based on BH3-only gene expression, grouped together the CGNP clusters that were enriched in *Eed^cKO^* mice (red box). **(B)** Heat map of BH3-only gene expression in the combined set of CGNP/CGN clusters, comparing *Eed^cKO^* and control genotypes. **(C)** Representative H&E-stained sagittal sections of P21 WT, *Eed^cKO^*, *Eed/Trp53^dKO^*, and *Eed/Bax/Bak^tKO^*mice. * indicates regions of IGL. Arrow indicates regions of persistent or absent EGL.

Hierarchical clustering based on BH3-only gene expression distinguished the 3 proliferative clusters enriched in *Eed^cKO^* cerebella, Clusters 4_1, 5, and 6_1, which showed increased expression of *Pmaip1* (aka *Noxa*), *Bad*, *Bcl2l11* (aka *Bim*), and *Hrk* (Fig. 4A). Similarly, comparing the combined set of all cells in CGNP and CGN clusters, *Eed^cKO^* cells showed a distinctive pattern of BH3-only gene expression, with increased *Bbc3* (aka *Puma*) *Pmaip1* (aka *Noxa*), *Bad*, *Bcl2l11* (aka *Bim*), and *Hrk* (Fig. 4B). These data suggested that *Eed* deletion may increase apoptosis by direct activation of the intrinsic apoptotic pathway.

To determine whether p53 signaling or the intrinsic apoptotic pathway contributed to cerebellar hypoplasia in *Eed^cKO^* mice, we combined *Eed* deletion with deletion of either *Trp53* or both *Bax* and *Bak*. We bred *Eed^cKO^* mice with *Trp53^fl/fl^*mice to generate *Math1-Cre/Eed^fl/fl^/Trp53^fl/fl^* (*Eed/Trp53^dKO^*) mice and bred *Eed^cKO^* mice with *Bax^fl/fl^/Bak^-/-^*mice to generate *Math1-Cre/Eed^fl/fl^/Bax^fl/fl^/Bak^-/-^*(*Eed/Bax/Bak^tKO^*) mice. *Eed/Trp53^dKO^* mice showed cerebellar hypoplasia similar to *Eed^cKO^* mice (Fig. 4C). In contrast, cerebella in *Eed/Bax/Bak^tKO^* mice were markedly less abnormal, with relatively increased CGNs within the IGL and more appropriate layering of Purkinje cells between the IGL and molecular layers (Fig. 4C). *Bax*/*Bak* co-deletion did not fully rescue the effects of *Eed* knockout, as the IGL remained less densely populated than WT controls and the molecular layer contained ectopic cells (Fig. 4C). The absence of rescue in *Eed/Trp53^dKO^* mice indicates that growth failure in the *Eed^cKO^* cerebella was p53-independent. The partial rescue by co-deletion of *Bax* and *Bak*, however, demonstrates that p53-independent activation of the intrinsic apoptosis pathway contributed to cerebellar hypoplasia in *Eed^cKO^* mice. *Bax*/*Bak* co-deletion also increased the myoid population (Supplemental Fig. 2), indicating that myoid cells were typcially removed from *Eed^cKO^* cerebella by apoptosis.

### PRC2 function is not required for SHH medulloblastoma tumorigenesis

Mutations that disrupt cerebellar growth may identify genes required for growth of medulloblastoma (49) and the PRC2 has been proposed as a target for medulloblastoma therapy. Therefore, to determine whether medulloblastomas, like CGNPs, depend on *Eed* and the PRC2, we bred *Eed^cKO^* and *Ezh2^cKO^* mice with *SmoM2* mice (50). *SmoM2* mice harbor a *Cre*-conditional transgene with an oncogenic allele of the SHH receptor component *Smo*. Mice that inherit both *Math1-Cre* and *SmoM2* develop medulloblastoma with 100% penetrance and without treatment die of tumor progression by P50 (6). We have shown that these tumors recapitulate the gene expression patterns and cellular diversity of SHH medulloblastomas resected from patients (3, 36, 51). By interbreeding *Eed^cKO^* and *Ezh2^cKO^*mice with *SmoM2* mice, we generated pups with the genotypes *Math1-Cre/Eed^fl/fl^/SmoM2* (*M-Smo/Eed^cKO^*) and *Math1-Cre/Ezh2^fl/fl^/SmoM2* (*M-Smo/Ezh2^cKO^*). To generate control mice with SHH medulloblastomas with intact PRC2, we bred *Math1-Cre* and *SmoM2* mice to generate *Math1-Cre/SmoM2* (*M-Smo*) controls. We then compared medulloblastomas in *M-Smo/Eed^cKO^*, *M-Smo/Ezh2^cKO^*, and *M-Smo* mice.

*M-Smo/Eed^cKO^, M-Smo/Ezh2^cKO^*, and *M-Smo* mice all developed medulloblastomas with 100% frequency by P10 (Fig. 5A). *M-Smo* tumors showed heterogeneous H3K27me3, with strongest H3K27me3 expression in the most differentiated elements, similar to WT cerebella (Fig. 5B). Deletion of either *Eed* or *Ezh2* effectively blocked PRC2 function, as both *M-Smo/Eed^cKO^* and *M-Smo/Ezh2^cKO^*showed no detectable H3K27me3 in tumor cells (Fig. 5B). *M-Smo/Eed^cKO^* and *M-Smo/Ezh2^cKO^* mice showed shorter survival times compared to controls, indicating that PRC2-mutant medulloblastomas progressed more quickly (Fig. 5C). Thus SHH medulloblastomas with PRC2 disruption did not show growth impairment similar to *Eed^cKO^*CGNPs.

**Figure 5.**
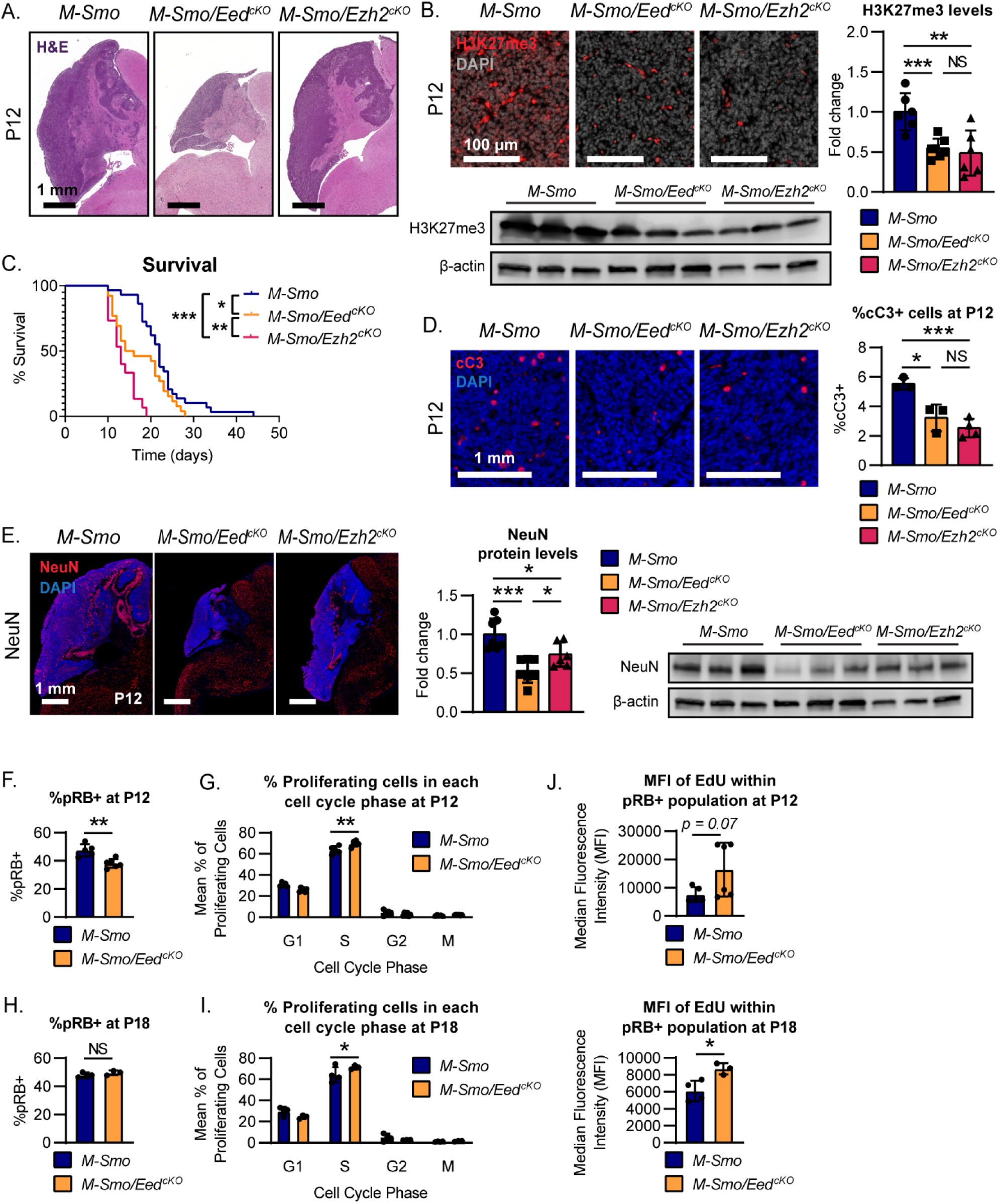
Loss of PCR2 function accelerated progression of SHH medulloblastomas. **(A)** Representative H&E-stained sagittal sections of medulloblastomas in *M-Smo*, *M-Smo/Eed^cKO^*, and *M-Smo/Ezh2^cKO^* mice at postnatal day 12 (P12). **(B)** Representative IF stains showing H3K27me3 in *M-Smo*, *M-Smo/Eed^cKO^*, and *MSmo/Ezh2^cKO^*tumors, and H3K27me3 western blots of medulloblastomas from 3 replicate mice of each genotype. **(C)** Kaplan-Meier curves compare the survival times of *M-Smo*, *M-Smo/Eed^cKO^*, and *MSmo/Ezh2^cKO^*mice. **(D)** IHC for cC3 in representative sections, with quantitative analysis of replicate samples. **(E)** IHC for NEUN in representative sections, with western blot on replicate samples quantified on the right. **(F-J)** flow cytometry analysis of dissociated tumors of indicated age and genotype, showing **(F, H)** pRB+ fractions and **(G,I)** cell cycle distribution of pRB+ cells, and **(J)** EdU MFI. *, **, and *** denote p<0.05, p<0.01 and p<0.001 respectively, relative to controls.

We compared apoptosis and terminal differentiation in *M-Smo/Eed^cKO^*, *M-Smo/Ezh2^cKO^*, and *M-Smo*, as these processes are important determinants of tumor growth. *Eed*-deleted and *Ezh2*-deleted medulloblastomas show smaller fractions of cC3+ cells compared to control tumors (Fig. 5D), indicating less apoptosis. PRC2-mutant medulloblastomas thus did not show the increased cell death that we noted in *Eed^cKO^* cerebella. Medulloblastomas in *M-Smo/Eed^cKO^* and *M-Smo/Ezh2^cKO^*mice also showed reduced neuronal differentiation, demonstrated by less abundance of neuronal marker NEUN (Fig. 5E). PRC2 disruption therefore did not increase apoptosis or terminal differentiation in SHH medulloblastoma.

### Transient growth suppression in *M-Smo/Eed^cKO^* medulloblastomas

While *M-Smo/Eed^cKO^* tumors progressed more rapidly than *M-Smo* control tumors, we noted that at P12, *M-Smo/Eed^cKO^*tumors were consistently smaller (Fig. 5A), suggesting that *Eed* deletion might produce an initial growth suppression, followed by more rapid growth. To analyze tumor growth dynamics, we compared RB phosphorylation and cell cycle progression in *M-Smo/Eed^cKO^* and *M-Smo* tumors. We injected EdU into 3-5 replicate mice of each genotype at either P12 or P18, then harvested tumors 1 hour after EdU injection and quantified pRB and EdU uptake by flow cytometry.

P12 *M-Smo/Eed^cKO^* tumors showed smaller fractions of pRB+ cells compared to P12 *M-Smo* control tumors (Fig. 5F), indicating that fewer tumor cells were proliferative. Within the pRB+ fractions of P12 *M-Smo/Eed^cKO^* tumors, however, more cells were in S-phase compared to the pRB+ fractions of P12 *M-Smo* tumors (Fig. 5G), indicating more rapid progression from G_1_. P12 *M-Smo/Eed^cKO^* tumors thus contained smaller proliferative populations, consistent with smaller tumor size. However, the cells that were proliferating in P12 *M-Smo/Eed^cKO^*medulloblastomas were cycling more rapidly, suggesting a transition to faster tumor growth.

Consistent with more rapid tumor growth after P12, by P18 *M-Smo/Eed^cKO^* tumors no longer showed fewer pRB+ cells (Fig. 5H). Moreover, pRB+ cells in *M-Smo/Eed^cKO^* tumors continued to show more rapid cycling, demonstrated by greater S-phase fractions, compared to pRB+ cells from P18 *M-Smo* tumors (Fig. 5I). Supporting the more rapid S-phase progression in *M-Smo/Eed^cKO^* tumors at both P12 and P18, pRB+ cells from *M-Smo/Eed^cKO^* tumors showed higher EdU median fluorescence intensity (MFI), indicating increased EdU uptake within the period of EdU exposure (Fig. 5J). The smaller pRB+ population in *M-Smo/Eed^cKO^* tumors at P12, the increased rate of proliferation within the pRB+ population, and the similar pRB+ population at P18 are all consistent with a biphasic effect of PRC2 disruption on tumor growth, in which tumor growth was initially decreased and then rebounded to produce shorter survival times.

### Up-regulation of PRC2 target genes without growth suppression in PRC2-mutant medulloblastomas

We investigated whether medulloblastomas with deletion of *Eed* or *Ezh2* up-regulated the same PRC2 targets that were up-regulated in *Eed^cKO^*CGNPs. Both *M-Smo/Eed^cKO^* and *M-Smo/Ezh2^cKO^*medulloblastomas showed frequent CDKN2A+ cells which were not observed in *M-Smo* control tumors (Fig. 6A). CDKN2A suppressed RB phosphorylation less effectively in tumor cells than in CGNPs, as the pRB+ fractions of CDKN2A+ cells were significantly higher in *M-Smo/Eed^cKO^* and *M-Smo/Ezh2^cKO^* medulloblastomas compared to P7 *Eed^cKO^*cerebella (Fig. 6B). Moreover, the pRB+ fraction of CDKN2A+ cells correlated with tumor growth; *M-Smo/Ezh2^cKO^*tumors, which progressed faster than *M-Smo/Eed^cKO^* tumors, showed higher pRB+ fractions of CDKN2A+ cells at P12. By P18, when proliferation accelerated in *M-Smo/Eed^cKO^* tumors, the pRB+ fractions of CDKN2A+ cells was also increased, to become similar to the P12 *M-Smo/Ezh2^cKO^*tumors. CDKN2A was thus up-regulated in PRC2-mutant medulloblastomas but did not restrict RB phosphorylation or tumor progression.

**Figure 6.**
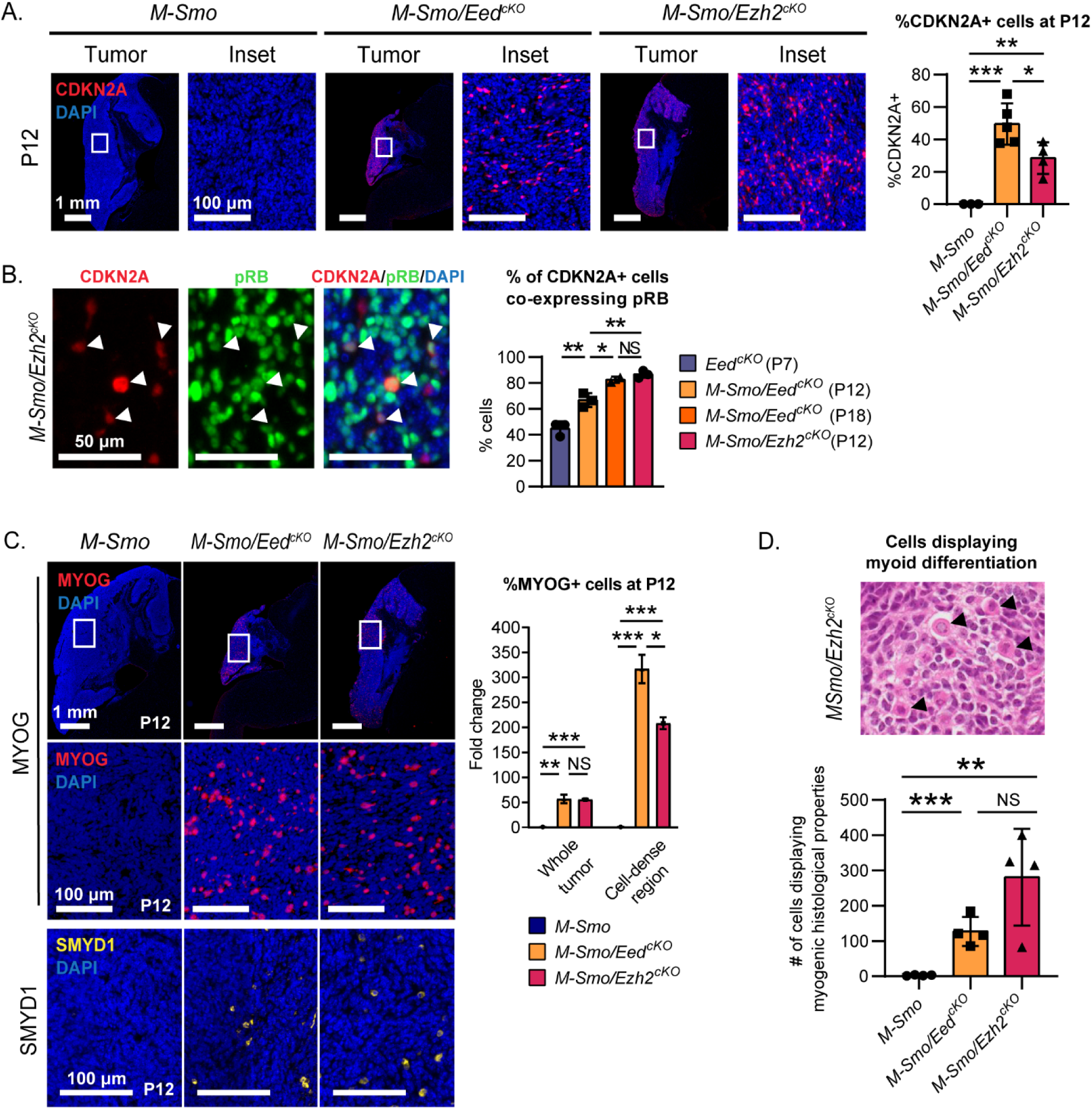
Up-regulation of PRC2 targets and myoid differentiation in both *Eed*-deleted and *Ezh2-*deleted SHH medulloblastomas. **(A)** Representative CDKN2A IHC in medulloblastomas of indicated genotypes, with quantification in replicate mice of each genotype. **(B)** Representative images of CDKN2A/pRB dual staining and quantification of pRB+ fractions of CDKN2A-expressing cells in the indicated ages and genotypes. **(C)** Representative MYOG and SMYD1 IHC in medulloblastomas of indicated genotypes with quantification of MYOG+ cells in replicate mice of each genotype. **(D)** Representative H&E-stained section from a medulloblastoma in a *M-Smo/Ezh2^cKO^* mouse, with myoid cells (arrows), and quantification in replicate mice of the indicated genotypes. *, **, and *** denote p<0.05, p<0.01 and p<0.001 respectively, relative to controls.

Analysis of muscle markers MYOG and SMYD1 showed inappropriate myoid differentiation in *M-Smo/Eed^cKO^* and *M-Smo/Ezh2^cKO^*tumors (Fig. 6C), as seen in *Eed^cKO^* cerebella. The expression of muscle genes in *M-Smo/Eed^cKO^* and *M-Smo/Ezh2^cKO^* medulloblastomas suggested similarity to the clinically observed medulloblastoma variant, medullomyoblastoma. To determine if the *M-Smo/Eed^cKO^* and *M-Smo/Ezh2^cKO^* tumors recapitulated medullomyoblastoma histopathology, we submitted H&E sections from replicate *M-Smo/Eed^cKO^, M-Smo/Ezh2^cKO^*, and *M-Smo* tumors to a blinded analysis. Two experienced pediatric neuropathologists counted myoid cells, defined by key morphologic changes (Fig. 6D), in replicate sections, while blinded to the genotype. The resulting quantifications of myoid cells correctly distinguished control tumors from tumors with either *Eed* or *Ezh2* deletion, which were not significantly different from each other (Fig. 6E). Deletion of either *Eed* or *Ezh2* was therefore sufficient to allow myoid differentiation, reproducing the molecular and histologic features of medullomyoblastoma.

## Discussion

Our data show that the PRC2 maintains neuronal fate commitment in cerebellar progenitors and in SHH medulloblastoma by preventing alternative, myoid differentiation. In the postnatal cerebellum, conditional deletion of PRC2 components *Ezh2* and *Eed* in the *Atoh1* lineage disrupted PRC2-mediated H3K27 trimethylation and caused a fraction of CGNPs to differentiate along a muscle cell trajectory. *Eed* deletion induced myoid differentiation in more cells than *Ezh2* deletion and markedly impaired cerebellar growth through a combination of decreased proliferation and increased apoptosis. In medulloblastomas, deletion of either *Eed* or *Ezh2* resulted in myoid differentiation, but neither deletion increased apoptosis or inhibited overall growth. Rather, deletion of either *Eed* or *Ezh2* accelerated tumor progression.

Single-cell transcriptomic analysis of postnatal cerebella showed that *Eed*-deleted CGNPs inappropriately expressed genes typically suppressed by the PRC2, including *Cdkn2a, Hoxa9*, and *Hoxa7*. Up-regulation of *Cdkn2a* may be sufficient to explain reduced CGNP proliferation, as seen in *Eed*-deleted hippocampal progenitors (11). Up-regulation of specific BH3-only genes suggested that direct activation of BAX or BAK may mediate increased CGNP apoptosis. Co-deletion studies of *Eed* plus either *Trp53* or *Bax* AND *Bak* confirmed that increased apoptosis occurred by p53-independent activation of the intrinsic apoptosis pathway. The partial rescue of cerebellar growth in *Eed/Bax/Bak^tKO^*mice demonstrates that inappropriate apoptosis contributed to growth failure and restricted the myoid population.

In *Eed*-deleted cerebella, most CGNPs were able to complete neural differentiation, as shown by the neural fates of Clusters 4_1 and 8 in *Eed^cKO^*mice. The inappropriate expression of *Hox* genes in these clusters did not prevent a recognizable CGN-like pattern of gene expression. In contrast, the myoid differentiation of Cluster 15 demonstrates that PRC2 disruption permitted new fate possibilities. These new fate possibilities were realized by a fraction CGNPs which no longer adhered to neural commitment. Based on the patterns of differentiation and gene expression, we propose that the PRC2 supports neural development by suppressing inappropriate transcriptional regulators such as *Myog*, and thus blocking progenitors from following inappropriate differentiation pathways.

*Eed* deletion resulted in cerebellar hypoplasia that was not seen in *Ezh2*-deleted mice. These discordant phenotypes may be explained by the distinct roles of each protein in the PRC2 complex. EED, which binds to trimethyl-lysine residues in histone tails, is required for PRC2 methyltransferase activity (52). In contrast, EZH2, which methylates lysine residues, can be partially compensated by the homolog EZH1 in multiple cellular contexts (53, 54). Compensation by EZH1 may allow sufficient PRC2 function to sustain cerebellar growth in *Ezh2^cKO^* mice, and to suppress PRC2 target gene expression in most but not all CGNPs, resulting in fewer myoid cells in *Ezh2^cKO^* cerebella compared to *Eed^cKO^*cerebella. Alternatively, as a shared component of the PRC1 and PRC2 complexes, EED loss may more broadly affect chromatin repression (55). However, our finding that *Eed* deletion did not affect levels of H2AK119 monoubiquitylation suggests that PRC1 activity was not altered in *Eed^cKO^* CGNPs or medulloblastomas. The overlapping patterns of differential gene expression in *Eed^cKO^* and *Ezh2^cKO^*cerebella provide evidence that *Eed* deletion and *Ezh2* deletion acted in CGNPs on the same regulatory pathway, with differences in number of cells affected driving the differences in phenotype.

Different phenotypes were similarly noted when either either *Eed* or *Ezh2* were deleted in intestinal epithelia (42). Conditional deletion of *Eed* in the intestinal crypts decreased proliferation and caused hypoplasia, while *Ezh2* deletion did not cause an overt phenotype. The continued proliferation in *Ezh2*-deleted intestinal crypts suggests that EZH1 may compensate for EZH2 loss in these cells (42), and a similar mechanism may explain sustained proliferation of *Ezh2-*deleted CGNPs.

In SHH medulloblastomas, disrupting PRC2 activity through deletion of either *Eed* or *Ezh2* was sufficient to allow widespread expression of genes typically suppressed by the PRC2, including the CDKN2A tumor suppressor. However, neither PRC2 disruption nor CDKN2A expression was sufficient for sustained suppression of tumor growth. *Eed* deletion reduced the average proliferation rate in P12 tumors, producing transient growth suppression. Over time, however, a fraction of *Eed*-deleted tumor cells that were rapidly proliferative increased, driving ultimately faster progression. The initial reduction in tumor growth in *M-Smo/Eed^cKO^* tumors was consistent with previous studies that showed anti-tumor effects of inhibiting EZH2 in SHH medulloblastoma (22, 56, 57). However, the shorter survival times in *M-Smo/Eed^cKO^* and *M-Smo/Ezh2^cKO^*mice raise concern that PRC2 disruption may not produce durable anti-tumor effects.

Our genetic studies identify a role of PRC2 in regulating neural fate commitment of cerebellar progenitors and medulloblastoma cells, and implicate PRC2 disruption in the pathogenesis of medullomyoblastoma, a subtype of medulloblastomas characterized by myogenic differentiation of tumor cells (58, 59). Our data also caution that pharmacologically inhibiting PRC2 function in medulloblastoma may hasten, rather than slow, tumor growth.

## Methods and Materials

### Mice

We generated *Ezh2^cKO^* mice by breeding *Math1-Cre* mice (Jackson Labs Stock #011104), which express Cre recombinase in CGNPs, with *Ezh2^LoxP/LoxP^* (Jackson Labs Stock #022616). We generated *Eed^cKO^* mice by breeding *Math1-Cre* mice with *Eed^LoxP/LoxP^* mice (generously donated by Dr. Terry Magnuson). To generate *Eed/Tp53^dKO^* mice, we interbred *Math1-Cre/Eed^LoxP/LoxP^*and *Trp53^LoxP/LoxP^* mice (Jackson Labs stock # 008462). To generate *Eed/Bax/Bak^tKO^* mice, we interbred *Math1-Cre/Eed^LoxP/LoxP^* and *Bax^LoxP/LoxP^/Bak^-/-^*mice (Jackson Labs stock # 006329).

To generate *M-Smo* mice, we crossed *Math1-Cre* mice with *SmoM2* mice (Jackson Labs stock #005131) that harbor a Cre-conditional transgene comprising of an oncogenic allele of *Smo*, fused to the YFP coding sequence. We then crossed *Math1-Cre/Ezh2^LoxP/LoxP^*and *Ezh2^LoxP/LoxP^/SmoM2^LoxP/LoxP^* mice to produce *M-Smo/Ezh2^cKO^* mice and *Math1-Cre/Eed^LoxP/LoxP^*and *Eed^LoxP/LoxP^/SmoM2^LoxP/LoxP^* mice to produce *M-Smo/Eed^cKO^* mice.

All mice were of species *Mus musculus* and crossed into the C57BL/6 background through at least five generations. All animal studies were carried out with the approval of the University of North Carolina Institutional Animal Care and Use Committee under IACUC protocols (19-098 and 21-011).

### Histology and Immunohistochemistry

Mouse brains were processed, immunostained, and quantified as previously described (Ocasio et al., 2019a; Malawsky et al., 2021; Lim et al., 2022; Lang et al., 2016). In brief, mice were placed under isoflurane anesthesia and decapitated. Harvested brains were fixed by immersion in 4% formaldehyde for 24 hours and then transferred to a graded ethanol series and embedded in paraffin and sectioned along the sagittal midline. Samples were stained and imaged using an Aperio Scanscope and quantified via automated cell counting using Tissue Studio (Definiens).

Primary antibodies used were: H3K27me3 diluted 1:200 (Cell Signaling, #9733), pRB diluted 1:3000 (Cell Signaling, #8516), cC3 diluted 1:400 (Biocare Medical, #CP229C), NeuN diluted 1:10000 (Millipore, MAB377), Myogenin (MYOG) diluted 1:500 (Abcam, ab124800), SMYD1 diluted 1:100 (ThermoFisher, PA5-84544), and CDKN2A diluted 1:500 (Abcam, ab241543). Stained images were counterstained with DAPI.

### Western Blot

Whole cerebella or tumors were harvested and homogenized in an SDS lysis buffer, which included SDS Solution (20%) (ThermoFisher 151-21-3), UltraPure™ 1 M Tris-HCI Buffer, pH 7.5 (ThermoFisher 15567027), UltraPure™ 0.5M EDTA, pH 8.0 (ThermoFisher 15575038), Phenylmethanesulfonyl fluoride (≥98.5%; powder) (Millipore Sigma 329-98-6), Isopropyl alcohol (Mallinckrodt 3037), and purified water. Equal total protein concentrations were loaded from each sample and run on SDS-polyacrylamide gels (BioRad, #4561105, #4568094), transferred onto polyvinylidene difluoride membranes, and membranes blotted using a SNAP i.d. 2.0 Protein Detection System (Millipore). The following antibodies were used: H3K27me3 (Cell Signaling, #9733; 1:500 dilution), H3K4me3 (Cell Signaling, #9751; 1:500 dilution), H2AK119ub (Cell Signaling, #8240; 1:500 dilution), H3K27Ac (Cell Signaling, #8173; 1:500 dilution), NeuN (Millipore, MAB377; 1:500 dilution), β-actin (Cell Signaling, #3700; 1:5000 dilution), Anti-rabbit IgG, HRP-linked antibody (Cell Signaling, #7074; 1.5:1000 dilution), and Anti-mouse IgG, HRP-linked antibody (Cell Signaling, #7076; 1.5:1000 dilution). Membranes were imaged using a chemiluminescent SuperSignal West Femto Maximum Sensitivity Substrate (34095, Thermo Fisher Scientific) and the C-DiGit blot scanner (LI-COR Biosciences). Blots were then quantified using Image Studio Lite software (LI-COR).

### Flow Cytometry – Cell cycle analysis

P12 and P18 *M-Smo* and *M-Smo/Eed^cKO^* mice were injected with Edu 1 hour prior to harvest. Mice were anesthetized with isoflurane and decapitated. Tumors samples were dissociated using the Cell Dissociation Kit (Worthington Biochemical Corporation, #LK003150), which included dissociation with papain at 37°C for 15min, and isolation using an ovomucoid inhibitor density gradient. Tumor cells were then treated with the Fixation and Permeabilization Kit (Life Technologies, #GAS004), and stained using the following antibodies: 647-conjugated pRB diluted 1:50 (Cell Signaling, #8974) and FxCycle Violet at 1:100 (Life Technologies, #F10347). Edu was detected using a Click-iT EdU Alexa Fluor 488 Imaging Kit (catalogue number C10337; Life Sciences). Samples were run on an LSRFortessa (BD Biosciences) at the UNC Flow Cytometry Core. Data was analyzed using Flow Jo v10.

### Single cell sequencing (scRNA-seq) – sample collection

Brains were harvested and cut along the sagittal midline. One half of the cerebellum from each mouse was dissociated and processed for scRNA-seq, and the other half of the brain was fixed, sectioned, and analyzed to confirm phenotype. Half cerebella were processed using the Cell Dissociation Kit (Worthington Biochemical Corporation, #LK003150), in which samples were treated with papain at 37°C for 15min and then separated by centrifugation of an ovomucoid inhibitor density gradient. Cells were then subjected to bead pairing by microfluidics, cDNA synthesis, and library construction using the Drop-seq V3 method (Macosko et al., 2015) as in our prior studies (Ocasio et al, 2019a).

### scRNA-seq – processing data

Data analysis was performed using the Seurat R package version 3.1.1 (Butler et al., 2018). Data were subjected to several filtering steps. Genes detected in > 30 cells were filtered out, to prevent misaligned reads appearing as rare transcripts in the data. Putative cells with fewer than 500 detected RNA molecules (nCount) or 200 different genes (nFeature) were considered to have too little information to be useful, and potentially to contain mostly ambient mRNA reads. Putative cells with greater than 4 standard deviations above the median nCount or nFeature were suspected to be doublets, improperly merged barcodes, or sequencing artifacts, and were excluded. As in our previously published work, putative cells with more than 10% mitochondrial transcripts were suspected to be dying cells and also excluded (Ocasio et al., 2019a).

In total, 86% of putative cells from WT mice and 74% of putative cells from *Eed^cKO^* mice met QC criteria and were included in the analysis. From the 5 WT mice, we included a total of 6558 cells with a range of 673–1852 cells per animal and a median of 1138 cells. From the 3 *Eed^cKO^*mice, we included a total of 2576 cells, with a range of 692–1036 cells per animal and a median of 847 cells.

### scRNA-seq – data normalization, clustering, differential gene expression, and cell type identification

The data was normalized using the SCTransform method as implemented in Seurat. The function then selected the top 3,000 most highly variable genes. PCA was performed on the subset of highly variable genes using the RunPCA function. We used 15 PCs in downstream analysis, based on examining the elbow in the elbow plot as implemented by Seurat. We identified cell clusters using the FindNeighbors and FindClusters functions.

To identify differential genes between clusters of cells, we used the Wilcoxon rank sum test to compare gene expression of cells within the cluster of interest to all cells outside that cluster, implemented by the FindMarkers function. Uniform Manifold Approximation and Projection (UMAP) was used to reduce the PCs to two dimensions for data visualization using the RunUMAP function. For re-iterated analysis of the Clusters 4 and 6, the same procedures were used. We then determined the type of cell within each cluster by analyzing cluster-specific gene expression patterns.

### Pathology Scoring

Sagittal H&E sections of P12 *M-Smo*, *M-Smo/Eed^cKO^*, and *M-Smo/Ezh2^cKO^* mouse brains were analyzed by neuropathologists (MS and JV) while blinded to the genotype, and the number of myoid cells per sample were manually counted.

### Survival Curves

Tumor-bearing mice were monitored daily and harvested according to a pre-determined humane endpoint, which included a decrease of weight >10% overnight, a hunched posture, decreased mobility or inability to eat, and ataxia.

### Statistical Analyses

Two-tailed Student’s t-tests were used to compare IHC and western blot quantifications between genotypes. A one-tailed 2 proportion z-test was used to compare MYOG+ cells between WT and *Ezh2^cKO^*cerebella. Survival curves were compared using the Log-rank (Manel-Cox) test. Dirichelet regression analysis was performed in R using the DirchletReg 0.7-1 package (60).

## Supplementary Data

Included in separate Excel document.

## Acknowledgments

We thank the UNC CGBID Histology Core supported by P30 DK 034987, and the UNC Tissue Pathology Laboratory Core supported by NCI CA016086 and UNC UCRF.

## Author contributions

Conceptualization: AHC, ID, TRG; Methodology: AHK, DM, TRG; Investigation: AHK, DM, MC, CR, FR; Visualization: AHC, DM, TRG; Funding acquisition: TRG; Supervision: TRG; Writing – original draft: AHK, DM, TRG; Writing – review & editing: AHK, DM, ID, TRG.

## Data availability

The scRNA-seq data are publicly available in the GEO database. The accession number for controls is GSE129730 and the accession number for *Eed^cKO^* is pending.

## Supplementary Figures

**Supplementary Figure 1.**
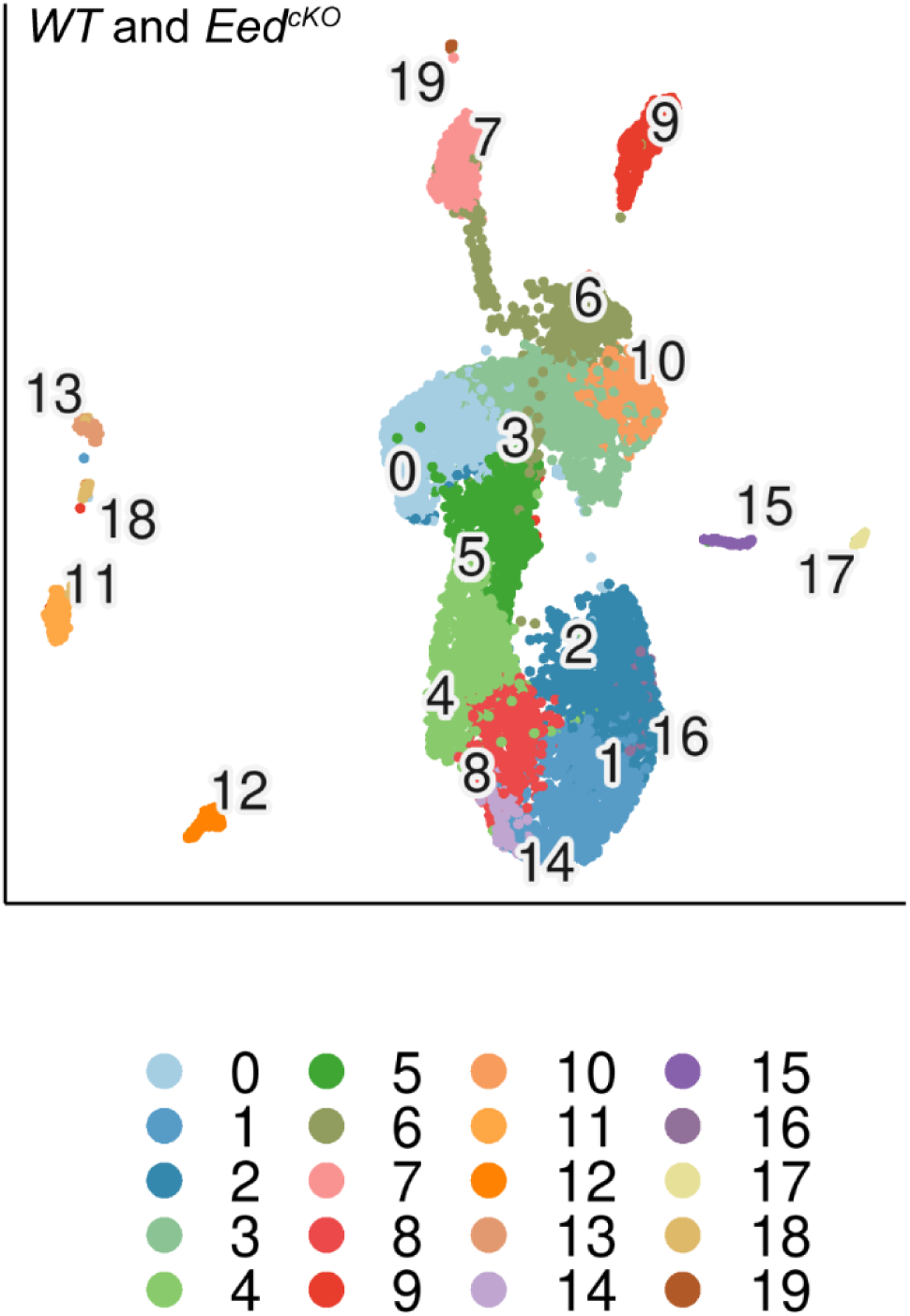
Initial uniform manifold approximation and projection (UMAP) qualitative map of cells dissociated from harvested cerebella from 5 WT and 3 *Eed^cKO^* mice. Cells were subdivided into 20 color-coded clusters.

**Supplemental Fig. 2:**
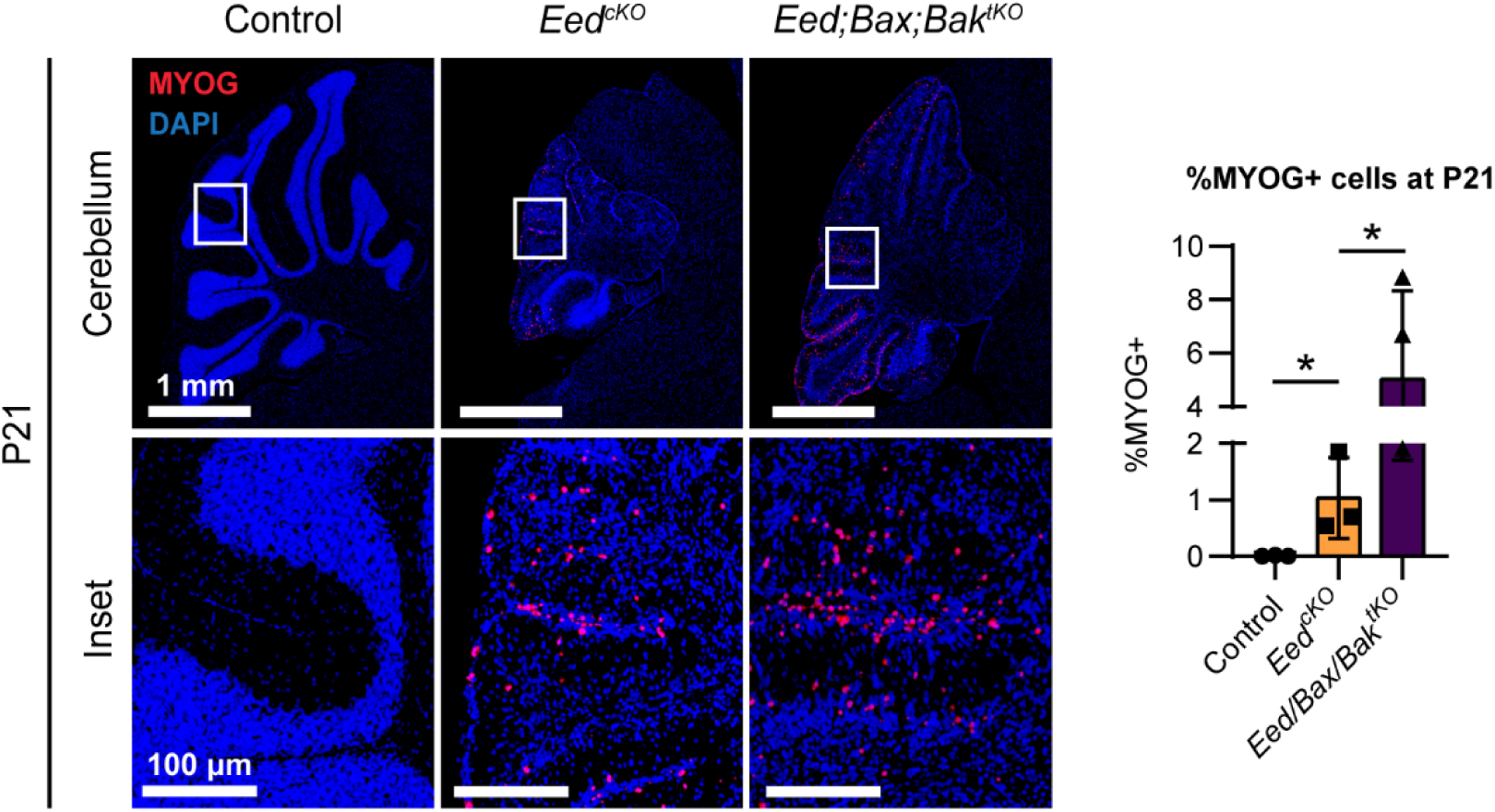
Disabling apoptosis increased the myoid population. Representative images of MYOG IHC in cerebella of indicated genotypes show that a population of myoid cells persisted in *Eed^cKO^*cerebella at P21, and that the myoid population increased when apoptosis was blocked by deletion of both *Bax* AND *Bak*. The increased myoid cells in *Eed/Bax/Bak^tKO^* cerebella indicates that apoptosis decreases the myoid population in *Eed^cKO^* cerebella.

## References

1. Roussel MF, Hatten ME. Cerebellum development and medulloblastoma. Curr Top Dev Biol. 2011;94:235–82. Epub 2011/02/08. doi: 10.1016/B978-0-12-380916-2.00008-5. PubMed PMID: 21295689; PMCID: 3213765.

2. Englund C, Kowalczyk T, Daza RA, Dagan A, Lau C, Rose MF, Hevner RF. Unipolar brush cells of the cerebellum are produced in the rhombic lip and migrate through developing white matter. J Neurosci. 2006;26(36):9184–95. Epub 2006/09/08. doi: 10.1523/JNEUROSCI.1610-06.2006. PubMed PMID: 16957075; PMCID: PMC6674506.

3. Ocasio J, Babcock B, Malawsky D, Weir SJ, Loo L, Simon JM, Zylka MJ, Hwang D, Dismuke T, Sokolsky M, Rosen EP, Vibhakar R, Zhang J, Saulnier O, Vladoiu M, El-Hamamy I, Stein LD, Taylor MD, Smith KS, Northcott PA, Colaneri A, Wilhelmsen K, Gershon TR. scRNA-seq in medulloblastoma shows cellular heterogeneity and lineage expansion support resistance to SHH inhibitor therapy. Nat Commun. 2019;10(1):5829. Epub 2019/12/22. doi: 10.1038/s41467-019-13657-6. PubMed PMID: 31863004; PMCID: PMC6925218.

4. Yang Z-J, Ellis T, Markant SL, Read T-A, Kessler JD, Bourboulas M, Schüller U, Machold R, Fishell G, Rowitch DH, Wainwright BJ, Wechsler-Reya RJ. Medulloblastoma Can Be Initiated by Deletion of Patched in Lineage-Restricted Progenitors or Stem Cells. Cancer Cell. 2008;14(2):135–45. doi: DOI: 10.1016/j.ccr.2008.07.003.

5. Oliver TG, Read TA, Kessler JD, Mehmeti A, Wells JF, Huynh TTT, Lin SM, Wechsler-Reya RJ. Loss of patched and disruption of granule cell development in a pre-neoplastic stage of medulloblastoma. Development. 2005;132(10):2425–39.

6. Schüller U, Heine VM, Mao J, Kho AT, Dillon AK, Han Y-G, Huillard E, Sun T, Ligon AH, Qian Y, Ma Q, Alvarez-Buylla A, McMahon AP, Rowitch DH, Ligon KL. Acquisition of Granule Neuron Precursor Identity Is a Critical Determinant of Progenitor Cell Competence to Form Shh-Induced Medulloblastoma. Cancer Cell. 2008;14(2):123–34. doi: DOI: 10.1016/j.ccr.2008.07.005.

7. Hallahan AR, Pritchard JI, Hansen S, Benson M, Stoeck J, Hatton BA, Russell TL, Ellenbogen RG, Bernstein ID, Beachy PA, Olson JM. The SmoA1 mouse model reveals that notch signaling is critical for the growth and survival of sonic hedgehog-induced medulloblastomas. Cancer Research. 2004;64(21):7794–800. Epub 2004/11/03. doi: 10.1158/0008-5472.CAN-04-1813. PubMed PMID: 15520185.

8. Hatten ME, Roussel MF. Development and cancer of the cerebellum. Trends in Neurosciences. 2011;34(3):134–42. Epub 2011/02/15. doi: 10.1016/j.tins.2011.01.002. PubMed PMID: 21315459; PMCID: 3051031.

9. Bailey P, Cushing, H. Medulloblastoma cerebelli: a common type of mid-cerebellar glioma of childhood. Arch Neurol Psychiatry. 1925;14:192–224.

10. Yao M, Ventura PB, Jiang Y, Rodriguez FJ, Wang L, Perry JSA, Yang Y, Wahl K, Crittenden RB, Bennett ML, Qi L, Gong CC, Li XN, Barres BA, Bender TP, Ravichandran KS, Janes KA, Eberhart CG, Zong H. Astrocytic trans-Differentiation Completes a Multicellular Paracrine Feedback Loop Required for Medulloblastoma Tumor Growth. Cell. 2020;180(3):502–20 e19. Epub 2020/01/28. doi: 10.1016/j.cell.2019.12.024. PubMed PMID: 31983537; PMCID: PMC7259679.

11. Liu PP, Xu YJ, Dai SK, Du HZ, Wang YY, Li XG, Teng ZQ, Liu CM. Polycomb Protein EED Regulates Neuronal Differentiation through Targeting SOX11 in Hippocampal Dentate Gyrus. Stem Cell Reports. 2019;13(1):115–31. Epub 2019/06/18. doi: 10.1016/j.stemcr.2019.05.010. PubMed PMID: 31204298; PMCID: PMC6627036.

12. Shi X, Zhang Z, Zhan X, Cao M, Satoh T, Akira S, Shpargel K, Magnuson T, Li Q, Wang R, Wang C, Ge K, Wu J. An epigenetic switch induced by Shh signalling regulates gene activation during development and medulloblastoma growth. Nat Commun. 2014;5:5425. Epub 2014/11/06. doi: 10.1038/ncomms6425. PubMed PMID: 25370275; PMCID: PMC5232137.

13. Wechsler-Reya RJ, Scott MP. Control of Neuronal Precursor Proliferation in the Cerebellum by Sonic Hedgehog. Neuron. 1999;22(1):103–14. doi: Doi: 10.1016/s0896-6273(00)80682-0.

14. Kenney AM, Rowitch DH. Sonic hedgehog Promotes G1 Cyclin Expression and Sustained Cell Cycle Progression in Mammalian Neuronal Precursors. Mol Cell Biol. 2000;20(23):9055–67. doi: 10.1128/mcb.20.23.9055-9067.2000.

15. Goodrich LV, Milenkovic L, Higgins KM, Scott MP. Altered neural cell fates and medulloblastoma in mouse patched mutants. Science. 1997;277(5329):1109–13. PubMed PMID: 9262482.

16. Taylor MD, Liu L, Raffel C, Hui CC, Mainprize TG, Zhang X, Agatep R, Chiappa S, Gao L, Lowrance A, Hao A, Goldstein AM, Stavrou T, Scherer SW, Dura WT, Wainwright B, Squire JA, Rutka JT, Hogg D. Mutations in SUFU predispose to medulloblastoma. Nat Genet. 2002;31(3):306–10. Epub 2002/06/18. doi: 10.1038/ng916. PubMed PMID: 12068298.

17. Vorechovsky I, Tingby O, Hartman M, Stromberg B, Nister M, Collins VP, Toftgard R. Somatic mutations in the human homologue of Drosophila patched in primitive neuroectodermal tumours. Oncogene. 1997;15(3):361–6. Epub 1997/07/17. doi: 10.1038/sj.onc.1201340. PubMed PMID: 9233770.

18. Wolter M, Reifenberger J, Sommer C, Ruzicka T, Reifenberger G. Mutations in the human homologue of the Drosophila segment polarity gene patched (PTCH) in sporadic basal cell carcinomas of the skin and primitive neuroectodermal tumors of the central nervous system. Cancer Res. 1997;57(13):2581–5. PubMed PMID: 9205058.

19. Pirrotta V, Gross DS. Epigenetic silencing mechanisms in budding yeast and fruit fly: different paths, same destinations. Mol Cell. 2005;18(4):395–8. Epub 2005/05/17. doi: 10.1016/j.molcel.2005.04.013. PubMed PMID: 15893722.

20. Obier N, Lin Q, Cauchy P, Hornich V, Zenke M, Becker M, Muller AM. Polycomb protein EED is required for silencing of pluripotency genes upon ESC differentiation. Stem Cell Rev Rep. 2015;11(1):50–61. Epub 2014/08/20. doi: 10.1007/s12015-014-9550-z. PubMed PMID: 25134795.

21. Yeo GW, Coufal N, Aigner S, Winner B, Scolnick JA, Marchetto MC, Muotri AR, Carson C, Gage FH. Multiple layers of molecular controls modulate self-renewal and neuronal lineage specification of embryonic stem cells. Hum Mol Genet. 2008;17(R1):R67–75. Epub 2008/07/18. doi: 10.1093/hmg/ddn065. PubMed PMID: 18632700.

22. Zhang H, Zhu D, Zhang Z, Kaluz S, Yu B, Devi NS, Olson JJ, Van Meir EG. EZH2 targeting reduces medulloblastoma growth through epigenetic reactivation of the BAI1/p53 tumor suppressor pathway. Oncogene. 2020;39(5):1041–8. Epub 2019/10/05. doi: 10.1038/s41388-019-1036-7. PubMed PMID: 31582835; PMCID: PMC7780546.

23. Huang J, Gou H, Yao J, Yi K, Jin Z, Matsuoka M, Zhao T. The noncanonical role of EZH2 in cancer. Cancer Sci. 2021;112(4):1376–82. Epub 2021/02/23. doi: 10.1111/cas.14840. PubMed PMID: 33615636; PMCID: PMC8019201.

24. Vo BT, Li C, Morgan MA, Theurillat I, Finkelstein D, Wright S, Hyle J, Smith SMC, Fan Y, Wang YD, Wu G, Orr BA, Northcott PA, Shilatifard A, Sherr CJ, Roussel MF. Inactivation of Ezh2 Upregulates Gfi1 and Drives Aggressive Myc-Driven Group 3 Medulloblastoma. Cell Rep. 2017;18(12):2907–17. Epub 2017/03/23. doi: 10.1016/j.celrep.2017.02.073. PubMed PMID: 28329683; PMCID: PMC5415387.

25. Scelfo A, Piunti A, Pasini D. The controversial role of the Polycomb group proteins in transcription and cancer: how much do we not understand Polycomb proteins? FEBS J. 2015;282(9):1703–22. Epub 2014/10/16. doi: 10.1111/febs.13112. PubMed PMID: 25315766.

26. Feng X, Juan AH, Wang HA, Ko KD, Zare H, Sartorelli V. Polycomb Ezh2 controls the fate of GABAergic neurons in the embryonic cerebellum. Development. 2016;143(11):1971–80. Epub 2016/04/14. doi: 10.1242/dev.132902. PubMed PMID: 27068104; PMCID: PMC4920161.

27. Wallace VA. Purkinje-cell-derived Sonic hedgehog regulates granule neuron precursor cell proliferation in the developing mouse cerebellum. Current Biology. 1999;9(8):445–8. doi: Doi: 10.1016/s0960-9822(99)80195-x.

28. Machold R, Fishell G. Math1 Is Expressed in Temporally Discrete Pools of Cerebellar Rhombic-Lip Neural Progenitors. Neuron. 2005;48(1):17–24. doi: DOI: 10.1016/j.neuron.2005.08.028.

29. Pasini D, Malatesta M, Jung HR, Walfridsson J, Willer A, Olsson L, Skotte J, Wutz A, Porse B, Jensen ON, Helin K. Characterization of an antagonistic switch between histone H3 lysine 27 methylation and acetylation in the transcriptional regulation of Polycomb group target genes. Nucleic Acids Res. 2010;38(15):4958–69. Epub 2010/04/14. doi: 10.1093/nar/gkq244. PubMed PMID: 20385584; PMCID: PMC2926606.

30. Lavarone E, Barbieri CM, Pasini D. Dissecting the role of H3K27 acetylation and methylation in PRC2 mediated control of cellular identity. Nat Commun. 2019;10(1):1679. Epub 2019/04/13. doi: 10.1038/s41467-019-09624-w. PubMed PMID: 30976011; PMCID: PMC6459869.

31. Dahmane N, Ruiz-i-Altaba A. Sonic hedgehog regulates the growth and patterning of the cerebellum. Development. 1999;126(14):3089–100.

32. Garcia I, Crowther AJ, Gama V, Miller CR, Deshmukh M, Gershon TR. Bax deficiency prolongs cerebellar neurogenesis, accelerates medulloblastoma formation and paradoxically increases both malignancy and differentiation. Oncogene. 2013;32(18):2304–14. doi: 10.1038/onc.2012.248. PubMed PMID: 22710714; PMCID: PMC3449008.

33. Veleta KA, Cleveland AH, Babcock BR, He YW, Hwang D, Sokolsky-Papkov M, Gershon TR. Antiapoptotic Bcl-2 family proteins BCL-xL and MCL-1 integrate neural progenitor survival and proliferation during postnatal cerebellar neurogenesis. Cell Death Differ. 2021;28(5):1579–92. Epub 2020/12/10. doi: 10.1038/s41418-020-00687-7. PubMed PMID: 33293647; PMCID: PMC8166857.

34. Luecken MD, Theis FJ. Current best practices in single-cell RNA-seq analysis: a tutorial. Mol Syst Biol. 2019;15(6):e8746. Epub 2019/06/21. doi: 10.15252/msb.20188746. PubMed PMID: 31217225; PMCID: PMC6582955.

35. Lim C, Dismuke T, Malawsky D, Ramsey JD, Hwang D, Godfrey VL, Kabanov AV, Gershon TR, Sokolsky-Papkov M. Enhancing CDK4/6 inhibitor therapy for medulloblastoma using nanoparticle delivery and scRNA-seq-guided combination with sapanisertib. Sci Adv. 2022;8(4):eabl5838. Epub 2022/01/27. doi: 10.1126/sciadv.abl5838. PubMed PMID: 35080986; PMCID: PMC8791615.

36. Malawsky DS, Weir SJ, Ocasio JK, Babcock B, Dismuke T, Cleveland AH, Donson AM, Vibhakar R, Wilhelmsen K, Gershon TR. Cryptic developmental events determine medulloblastoma radiosensitivity and cellular heterogeneity without altering transcriptomic profile. Commun Biol. 2021;4(1):616. Epub 2021/05/23. doi: 10.1038/s42003-021-02099-w. PubMed PMID: 34021242; PMCID: PMC8139976.

37. Rosenberg AB, Roco CM, Muscat RA, Kuchina A, Sample P, Yao Z, Graybuck LT, Peeler DJ, Mukherjee S, Chen W, Pun SH, Sellers DL, Tasic B, Seelig G. Single-cell profiling of the developing mouse brain and spinal cord with split-pool barcoding. Science. 2018;360(6385):176–82. Epub 2018/03/17. doi: 10.1126/science.aam8999. PubMed PMID: 29545511.

38. Malawsky D, Gershon T. scRNA-seq for microcephaly research [4]: Dirichlet regression for single cell population differences. Methods in Molecular Biology book series. 2022;in press.

39. Stojic L, Jasencakova Z, Prezioso C, Stutzer A, Bodega B, Pasini D, Klingberg R, Mozzetta C, Margueron R, Puri PL, Schwarzer D, Helin K, Fischle W, Orlando V. Chromatin regulated interchange between polycomb repressive complex 2 (PRC2)-Ezh2 and PRC2-Ezh1 complexes controls myogenin activation in skeletal muscle cells. Epigenetics Chromatin. 2011;4:16. Epub 2011/09/07. doi: 10.1186/1756-8935-4-16. PubMed PMID: 21892963; PMCID: PMC3180244.

40. Guo J, Cai J, Yu L, Tang H, Chen C, Wang Z. EZH2 regulates expression of p57 and contributes to progression of ovarian cancer in vitro and in vivo. Cancer Sci. 2011;102(3):530–9. Epub 2011/01/06. doi: 10.1111/j.1349-7006.2010.01836.x. PubMed PMID: 21205084.

41. Yang X, Karuturi RK, Sun F, Aau M, Yu K, Shao R, Miller LD, Tan PB, Yu Q. CDKN1C (p57) is a direct target of EZH2 and suppressed by multiple epigenetic mechanisms in breast cancer cells. PLoS One. 2009;4(4):e5011. Epub 2009/04/03. doi: 10.1371/journal.pone.0005011. PubMed PMID: 19340297; PMCID: PMC2659786.

42. Koppens MA, Bounova G, Gargiulo G, Tanger E, Janssen H, Cornelissen-Steijger P, Blom M, Song JY, Wessels LF, van Lohuizen M. Deletion of Polycomb Repressive Complex 2 From Mouse Intestine Causes Loss of Stem Cells. Gastroenterology. 2016;151(4):684–97 e12. Epub 2016/06/28. doi: 10.1053/j.gastro.2016.06.020. PubMed PMID: 27342214.

43. Jacobs JJ, Kieboom K, Marino S, DePinho RA, van Lohuizen M. The oncogene and Polycomb-group gene bmi-1 regulates cell proliferation and senescence through the ink4a locus. Nature. 1999;397(6715):164–8. Epub 1999/01/29. doi: 10.1038/16476. PubMed PMID: 9923679.

44. Jürgens G. A group of genes controlling the spatial expression of the bithorax complex in Drosophila. Nature. 1985;316(6024):153–5. doi: 10.1038/316153a0.

45. Nagel S, Venturini L, Marquez VE, Meyer C, Kaufmann M, Scherr M, MacLeod RA, Drexler HG. Polycomb repressor complex 2 regulates HOXA9 and HOXA10, activating ID2 in NK/T-cell lines. Mol Cancer. 2010;9:151. Epub 2010/06/23. doi: 10.1186/1476-4598-9-151. PubMed PMID: 20565746; PMCID: PMC2894765.

46. Lee Y, McKinnon PJ. ATM dependent apoptosis in the nervous system. Apoptosis. 2000;5(6):523–9. Epub 2001/04/17. PubMed PMID: 11303911.

47. Crowther AJ, Ocasio JK, Fang F, Meidinger J, Wu J, Deal AM, Chang SX, Yuan H, Schmid R, Davis I, Gershon TR. Radiation Sensitivity in a Preclinical Mouse Model of Medulloblastoma Relies on the Function of the Intrinsic Apoptotic Pathway. Cancer Res. 2016;76(11):3211–23. Epub 2016/05/20. doi: 10.1158/0008-5472.CAN-15-0025. PubMed PMID: 27197166; PMCID: PMC4891232.

48. Crowther AJ, Gama V, Bevilacqua A, Chang SX, Yuan H, Deshmukh M, Gershon TR. Tonic activation of Bax primes neural progenitors for rapid apoptosis through a mechanism preserved in medulloblastoma. The Journal of neuroscience: the official journal of the Society for Neuroscience. 2013;33(46):18098–108. Epub 2013/11/15. doi: 10.1523/JNEUROSCI.2602-13.2013. PubMed PMID: 24227720; PMCID: 3828463.

49. Lang PY, Gershon TR. A New Way to Treat Brain Tumors: Targeting Proteins Coded by Microcephaly Genes?: Brain tumors and microcephaly arise from opposing derangements regulating progenitor growth. Drivers of microcephaly could be attractive brain tumor targets. Bioessays. 2018;40(5):e1700243. Epub 2018/03/27. doi: 10.1002/bies.201700243. PubMed PMID: 29577351; PMCID: PMC5910257.

50. Mao J, Ligon KL, Rakhlin EY, Thayer SP, Bronson RT, Rowitch D, McMahon AP. A novel somatic mouse model to survey tumorigenic potential applied to the Hedgehog pathway. Cancer Res. 2006;66(20):10171–8. Epub 2006/10/19. doi: 10.1158/0008-5472.CAN-06-0657. PubMed PMID: 17047082; PMCID: PMC3806052.

51. Riemondy KA, Venkataraman S, Willard N, Nellan A, Sanford B, Griesinger AM, Amani V, Mitra S, Hankinson TC, Handler MH, Sill M, Ocasio J, Weir SJ, Malawsky DS, Gershon TR, Garancher A, Wechsler-Reya RJ, Hesselberth JR, Foreman NK, Donson AM, Vibhakar R. Neoplastic and Immune single cell transcriptomics define subgroup-specific intra-tumoral heterogeneity of childhood medulloblastoma. Neuro Oncol. 2021;noab135. doi: 10.1093/neuonc/noab135.

52. Margueron R, Justin N, Ohno K, Sharpe ML, Son J, Drury WJ, 3rd, Voigt P, Martin SR, Taylor WR, De Marco V, Pirrotta V, Reinberg D, Gamblin SJ. Role of the polycomb protein EED in the propagation of repressive histone marks. Nature. 2009;461(7265):762–7. Epub 2009/09/22. doi: 10.1038/nature08398. PubMed PMID: 19767730; PMCID: PMC3772642.

53. Yoo KH, Oh S, Kang K, Hensel T, Robinson GW, Hennighausen L. Loss of EZH2 results in precocious mammary gland development and activation of STAT5-dependent genes. Nucleic Acids Res. 2015;43(18):8774–89. Epub 2015/08/08. doi: 10.1093/nar/gkv776. PubMed PMID: 26250110; PMCID: PMC4605299.

54. Shen X, Liu Y, Hsu YJ, Fujiwara Y, Kim J, Mao X, Yuan GC, Orkin SH. EZH1 mediates methylation on histone H3 lysine 27 and complements EZH2 in maintaining stem cell identity and executing pluripotency. Mol Cell. 2008;32(4):491–502. Epub 2008/11/26. doi: 10.1016/j.molcel.2008.10.016. PubMed PMID: 19026780; PMCID: PMC2630502.

55. Cao Q, Wang X, Zhao M, Yang R, Malik R, Qiao Y, Poliakov A, Yocum AK, Li Y, Chen W, Cao X, Jiang X, Dahiya A, Harris C, Feng FY, Kalantry S, Qin ZS, Dhanasekaran SM, Chinnaiyan AM. The central role of EED in the orchestration of polycomb group complexes. Nat Commun. 2014;5:3127. Epub 2014/01/25. doi: 10.1038/ncomms4127. PubMed PMID: 24457600; PMCID: PMC4073494.

56. Cheng Y, Liao S, Xu G, Hu J, Guo D, Du F, Contreras A, Cai KQ, Peri S, Wang Y, Corney DC, Noronha AM, Chau LQ, Zhou G, Wiest DL, Bellacosa A, Wechsler-Reya RJ, Zhao Y, Yang ZJ. NeuroD1 Dictates Tumor Cell Differentiation in Medulloblastoma. Cell Rep. 2020;31(12):107782. Epub 2020/06/25. doi: 10.1016/j.celrep.2020.107782. PubMed PMID: 32579914; PMCID: PMC7357167.

57. Miele E, Valente S, Alfano V, Silvano M, Mellini P, Borovika D, Marrocco B, Po A, Besharat ZM, Catanzaro G, Battaglia G, Abballe L, Zwergel C, Stazi G, Milite C, Castellano S, Tafani M, Trapencieris P, Mai A, Ferretti E. The histone methyltransferase EZH2 as a druggable target in SHH medulloblastoma cancer stem cells. Oncotarget. 2017;8(40):68557–70. Epub 2017/10/06. doi: 10.18632/oncotarget.19782. PubMed PMID: 28978137; PMCID: PMC5620277.

58. Benson R, Mallick S, Bhanu Prasad V, Haresh KP, Gupta S, Julka PK, Rath GK. Medullomyoblastoma treated with craniospinal radiation and adjuvant chemotherapy: Report of 4 cases and review of the literature. J Egypt Natl Canc Inst. 2015;27(2):109–11. Epub 2015/05/06. doi: 10.1016/j.jnci.2015.03.001. PubMed PMID: 25936510.

59. Sachdeva MU, Vankalakunti M, Rangan A, Radotra BD, Chhabra R, Vasishta RK. The role of immunohistochemistry in medullomyoblastoma--a case series highlighting divergent differentiation. Diagn Pathol. 2008;3:18. Epub 2008/04/29. doi: 10.1186/1746-1596-3-18. PubMed PMID: 18439235; PMCID: PMC2391140.

60. Maier M. DirichletReg: Dirichlet Regression for Compositional Data in R. Research Report Series / Department of Statistics and Mathematics, WU Vienna University of Economics and Business, Vienna. 2014;125.

